# Cerebellar Bergmann Glia Integrating Noxious Information Modulate Nocifensive Behaviors

**DOI:** 10.1101/2022.05.18.489807

**Authors:** Seung Ha Kim, Jaegeon Lee, Seung-Eon Roh, Mirae Jang, Soobin Kim, Ji Hwan Lee, Jewoo Seo, Jae Yoon Hwang, Yong-Seok Lee, Eiji Shigetomi, C. Justin Lee, Schuichi Koizumi, Sun Kwang Kim, Sang Jeong Kim

**Affiliations:** Department of Physiology, Seoul National University College of Medicine, Seoul, Korea; Department of Biomedical Sciences, Seoul National University College of Medicine, Seoul, Korea; Department of Physiology, College of Korean Medicine, Kyung Hee University, Seoul, Korea; Memory Network Medical Research Center, Neuroscience Research Institute, Wide River Institute of Immunology, Seoul National University College of Medicine, Seoul, Korea; Department of Neuropharmacology, University of Yamanashi, Yamanashi, Japan; Yamanashi GLIA Center, Interdisciplinary Graduate School of Medicine, University of Yamanashi, Yamanashi, Japan; Center for Cognition and Sociality, Institute for Basic Science, Daejeon, Korea

## Abstract

Clinical studies have revealed that the cerebellum is activated by noxious stimuli or pathological pain, and its removal results in somatosensory dysfunction. However, the neural circuits and molecular mechanisms underlying the processing of noxious information in the cerebellum remain unknown. Using two-photon microscopy and optogenetics in mice, we found that the locus coeruleus (LC) terminals in the cerebellar cortex release noradrenaline (NA) in response to cutaneous noxious electrical stimuli. Most Bergmann glia (BG) accumulated this LC-NA noxious information by increasing intracellular calcium in an integrative manner. This global calcium activation of BG, referred to as “flare,” was also elicited in response to an intraplantar capsaicin injection. Chemogenetic inactivation of LC terminals or BG in the cerebellar cortex suppressed BG flares and reduced licking, a nocifensive behavior associated with capsaicin-induced pain. BG-specific knockdown of α-1 adrenergic receptors also suppressed capsaicin-induced BG flares and licking. Chemogenetic activation of BG or an intraplantar capsaicin injection reduced Purkinje cell firings, which disinhibited the output activity of the deep cerebellar nuclei. These results suggest that BG in the cerebellar cortex play an essential role in computing noxious information ascending from the LC and modulate pain-related behaviors by controlling the activity of the cerebellar neural circuits.

**One Sentence Summary:** Bergmann glia mediate noxious information processing in the cerebellum

## Introduction

Pain is defined as “an unpleasant sensory and emotional experience associated with or resembling that associated with actual or potential tissue damage” [1]. Thus, the processing of noxious information likely involves multiple regions of the brain [2, 3], among which the cerebellum has been relatively neglected in the field of pain medicine. The cerebellum plays an important role in coordinating movement while receiving diverse sensory inputs [4]. Interestingly, human brain imaging studies have consistently reported that the cerebellum is activated during physiological and pathological states of pain [5–7], while research on cerebellar stimulation has shown contradictory results (e.g., analgesia or hyperalgesia), depending on the method and site of stimulation [8]. To our knowledge, no study has revealed how peripheral noxious information is transmitted and processed within the cerebellar cortex.

The noradrenergic system transmits, processes, and modulates pain information throughout the brain [9]. The locus coeruleus (LC), a major source of noradrenaline (NA), is activated by noxious peripheral stimuli [10]. The LC has ascending projections to the prefrontal cortex and limbic system, which are known as pain-related higher brain centers, as well as the cerebellum [11]. Activated LC neurons release NA from their nerve terminals in the form of volume transmission, affecting the innervated brain regions [12, 13]. Previous studies have shown that the innervation of the cerebellum by the LC-NA system is actively involved in motor and cognitive behaviors [14, 15]. Therefore, we postulated that the cerebellum receives pain signals via the LC-NA system.

Astrocytes fulfill critical roles in the functional and structural changes of neuronal plasticity [16] and in disease conditions, such as pain processing, as well as in various physiological functions [17, 18] Bergmann glia (BG) (i.e., cerebellar cortical radial astrocytes) are closely associated with Purkinje cells (PCs) (i.e., cerebellar cortical sole output neurons) and can modulate their activity [19–21]. As radial astrocytes in zebrafish accumulate NA signals [22] and BG strongly respond to NA inputs by elevating intracellular calcium [14, 23, 24], LC-NA-activated BG is a strong candidate for the mediation of cerebellar pain processing.

In this study, we reveal that peripheral noxious stimuli induce the release of NA from LC terminals in the cerebellar cortex, triggering BG calcium activation. We also show that alpha-1 adrenergic receptors (α1-ARs) of BG are essential for LC-BG activation and nocifensive behavior, a tonic pain behavioral phenotype. Finally, we demonstrate that α1-AR-mediated BG activation changes the PC outputs during pain. Our findings provide a critical mechanism of the modulation of cerebellar noxious information, which could represent a novel therapeutic target for pain medicine.

## Result

To evaluate whether LC neurons project axons to the cerebellar cortex, we applied dual injections of AAV-TH-cre and AAV-DIO-mCherry to the LC to achieve mCherry expression in LC neurons that were co-labeled with dopamine beta-hydroxylase, an LC neuronal marker. Colabeled signals were also observed in the cerebellar cortex vermal lobule 4/5 (Fig. 1a), which is a part of the spinocerebellum that receives somatosensory input [25]. Next, to examine whether the LC transmits noxious information to the cerebellar cortex, we co-expressed the genetically encoded calcium indicator (GCaMP6s) and inhibitory DREADD (hM4Di) in LC neurons (Fig. 1b). *In vivo* two-photon imaging assisted by CNMF_E region-of-interest detection showed that the calcium level of LC terminals in the molecular layer of the cerebellar cortex increased in response to noxious electrical stimuli applied to the hind paw of anesthetized mice. The noxious stimuli-induced calcium signals of LC terminals were diminished when compound 21 (C21), an hM4Di agonist, was locally injected into the cerebellum (Fig. 1c–e). These results indicate that the LC conveys noxious information to the cerebellum.

**Fig.1:**
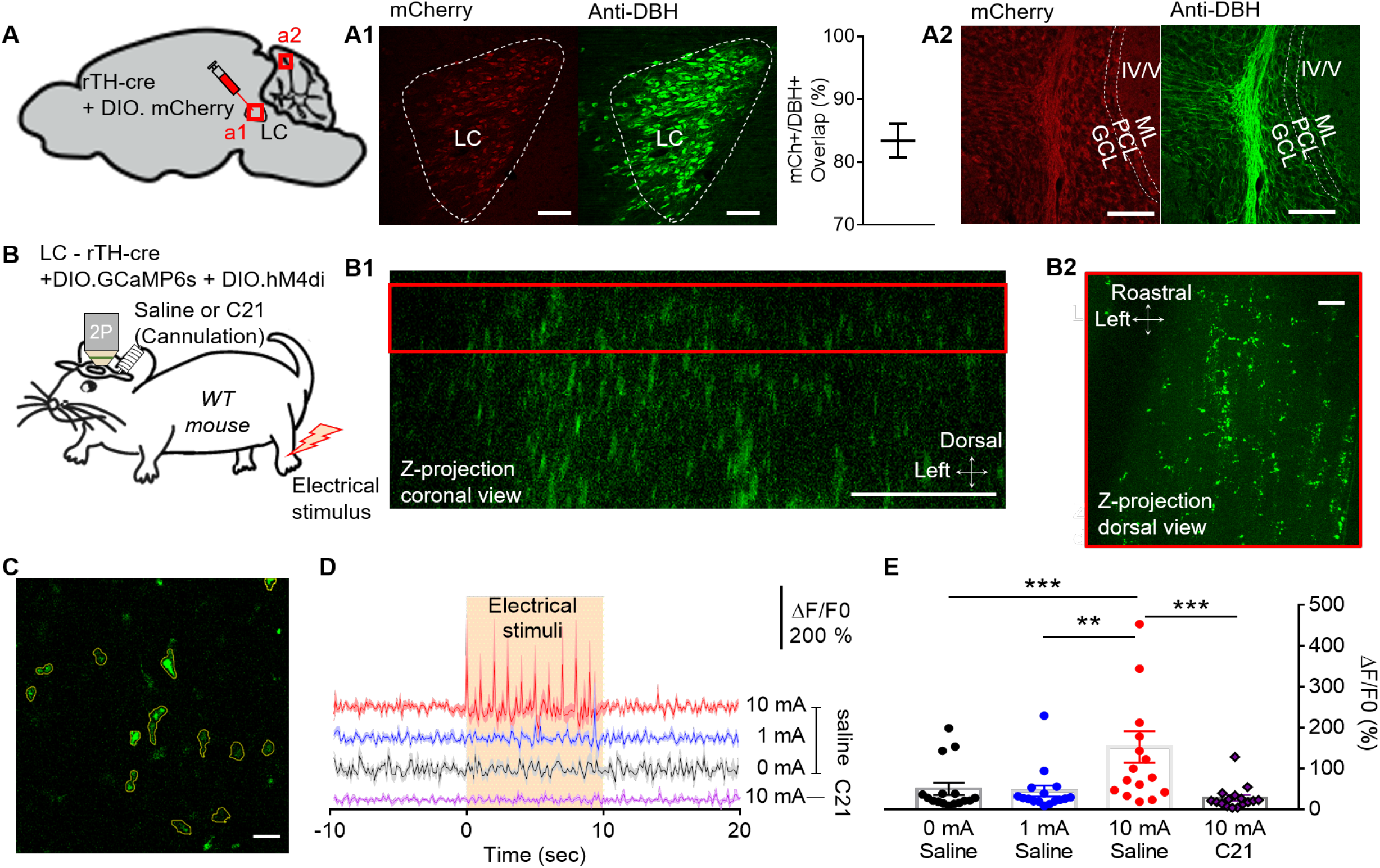
LC terminal activation in the cerebellar cortex by noxious stimuli. **A,** Schematic diagram showing virus injection for LC-specific mCherry expression. **A1-2**, mCherry was expressed in LC and cerebellar cortex. mCherry (red), DBH (dopamine beta hydroxylase, Green); Scale bars, 100 μm. Ratio of viral expression overlap with DBH positive cells in LC (83.39 ± 3.80 %; n = 2 slices of 1 mouse). **B,** Schematic diagram showing *in vivo* two-photon calcium imaging using hind-paw electrical in LC-specific hM4Di and GECI expressing mouse. **B1,**. A coronal view of the z-stack image (red box, molecular layer, 40 μm from the surface). **B2,** A dorsal view of the z-stack image, which represents the molecular layer (red box region of b1); Scale bars, 100 μm. **C,** An example of ROI detection of LC terminals using the CNMF_E. **D,** Representative traces of LC-terminal signals during various electrical stimuli (0, 1, 10 mA; 0.2 ms, 3 Hz, 10 sec) on the hindpaw (n = 16 ROIs in 3 mice). **E,** Increased peak of LC terminal calcium by hind-paw electrical stimuli and blocked by LC terminal hM4Di activation (n = 16 ROIs in 3 mice; 0 mA and 10 mA in saline group, *** p = 0.0004; 1 mA and 10 mA in saline group, ** p < 0.0013; 10 mA in saline group and 10 mA in c21 group, *** p = 0.0008; Wilcoxon test).

As BG activation has been reported to be NA-dependent [14], the LC may convey noxious information to the cerebellum via BG. To test how LC terminals transmit noxious information to BG, we performed simultaneous two-photon calcium imaging of LC terminals and BG in the molecular layer of the cerebellar cortex (Fig. 2a). Noxious electrical stimuli to the hind paw immediately increased the calcium signals in the LC terminals, whereas the BG calcium signal gradually increased to its peak amplitude (peak latency: 6.65 ± 0.17 s, Fig. 2b) with a delay of several seconds. Quantitative analysis showed that the calcium signals from the LC terminals and BG were both significantly increased after noxious stimulation (Fig. 2c).

**Fig.2:**
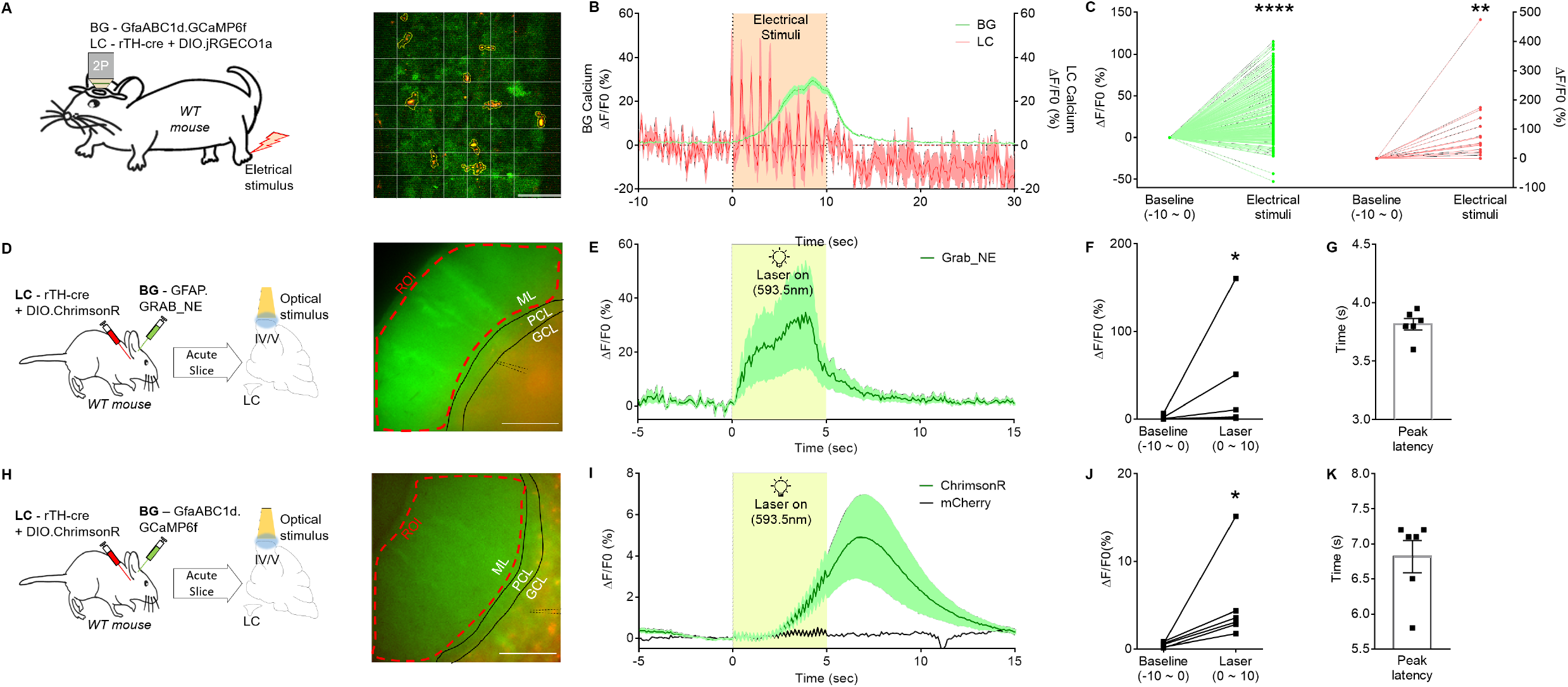
NA released from LC terminal activates cerebellar BG. **A,** Schematic diagram showing *in vivo* two-photon calcium imaging using hind-paw electrical stimulus in BG and LC GECI expressing mouse. An example of ROI detection of BG using grid pattern (8×8 ROIs) and LC terminal using the CNMF_E. Scale bar, 100 μm. **B,** Traces of LC terminal and BG calcium signal during hind-paw noxious stimuli (0.2 ms, 10 mA, 3 Hz; BG, n = 256 ROIs in 4 mice; LC terminal, n = 17 ROIs in 4 mice; Line for Mean, shade for SEM). **C,** Increased BG and LC terminal calcium activity by hind-paw electrical stimulation (BG, n = 256 ROIs in 4 mice, **** p < 0.0001; LC terminal, n = 17 ROIs in 4 mice, ** p = 0.0057; paired t test). **D,** Schematic diagram showing in-vivo virus injection for LC neuronal optogenetic modulation and BG Grab_NE imaging in cerebellar slice. An example of ROI detection of BG global activity in molecular layer; Scale bar, 100 μm. **E,** Trace of BG-specific NE sensor signal during LC optogenetic activation (593.5 nm, 5 mW, 5 Hz, 5 sec; n = 6 slices in 5 mice; Line for Mean, shade for SEM). **F,** Increased BG Grab_NE activity by laser stimulation (n = 6 slices; * P = 0.0313; Wilcoxon test). **G,** Latency of BG Grab_NE peak formation by laser stimulation (n = 6 slices; Mean ± SEM = 3.82 ± 0.05 s). **H,** Schematic diagram showing virus injection for LC neuronal optogenetic modulation and BG calcium slice imaging. An example of ROI detection of BG global activity in molecular layer; Scale bar, 100 μm. **I,** Trace of BG calcium signal during LC optogenetic activation or control (ChrimsonR, n = 6 slices in 5 mice; Control, n = 3 slices in 2 mice; Line for Mean, shade for SEM). **J,** Increased BG calcium activity by laser stimulation (n = 6 slices; * P = 0.313; Wilcoxon test). **K,** Latency of calcium peak formation by laser stimulation (n = 6 slices; Mean ± SEM = 6.82 ± 0.23 s)

Based on the order and temporal kinetics of calcium activation in the LC terminal and BG, we reasoned that LC terminals release NA upon stimulation, while BG accumulate the noxious information as glia in the fish accumulate evidence of futility [22]. To test this idea, we expressed an NA-sensor (Grab-NE) in BG [26] to measure the precise dynamics of NA acting on BG. At the same time, we expressed a red-shift opsin (ChrimsonR) in the LC to directly activate LC terminals in the cerebellar cortex (Fig. 2d, Fig. S1). The optogenetic activation of the LC terminal in the cerebellar slices revealed that NA levels acting on BG increased from the onset of laser stimulation, rapidly peaking before the last laser stimulus (peak latency: 3.82 ± 0.05 s) and returned to the baseline when the laser was turned off (Fig. 2e-g). Upon the same optogenetic activation of LC terminals, the BG calcium levels increased after a delay from the laser onset and continued to elevate even after the cessation of LC activation (peak latency: 6.82 ± 0.23 s) and slowly diminished thereafter (Fig. 2h-k). Finally, we tested whether BG integrate noxious information by increasing the number of electrical stimuli applied to the hind paw (Fig. S2). *In vivo* two-photon imaging revealed that BG calcium increased with increasing numbers of electrical stimuli. These results suggest that LC terminals in the cerebellar cortex transmit the noxious information to BG by releasing NA, and BG integrate this information.

We used a capsaicin-induced pain model to determine the role of the LC-NA-activated BG in the nocifensive behavior associated with tonic pain [27, 28]. Capsaicin activates TRPV1-positive nociceptors, which are required to elicit coping responses to tonic pain [28]. When capsaicin was injected into the sole of the hind paw, licking responses were evoked, likely to soothe the inevitable tonic pain (Fig. 3a). We used this nocifensive behavioral response to intraplantar capsaicin injection as a proxy to evaluate the degree of tonic pain. In a separate experiment, we measured BG calcium in response to the same intraplantar capsaicin injection using *in vivo* two-photon imaging. BG showed a distinct pattern of global calcium elevation, consistent with that observed from noxious electrical stimulation (Fig. 3b, c). This so-called “flare” is characterized by synchronized and simultaneous activation of the total BG [29]. In our 5-min recordings after capsaicin injection, BG displayed 2-5 flare activity (Fig. 3c). To evaluate whether the BG flare plays a role in nocifensive behavior, we inhibited this flare using a BG-specific chemogenetic approach. A previous study reported that hM4Di expressed in hippocampal CA1 astrocytes inhibited their calcium signals [30] (but also see [31]). Therefore, to inhibit the BG flare *in vivo*, we specifically infected BG with an AAV expressing hM4Di under the control of the astrocytic GFAP promoter. Additionally, we imaged GCaMP6f-expressing BG before and after intraperitoneal CNO injection. We first used noxious electrical stimulations that were available in a repetitively reproducible manner. BG calcium began to decrease 30 min after CNO administration in response to the noxious electrical stimulations, and the effect of inhibition was almost complete 1-2 hours after CNO administration (Fig. S3). Based on these data, we analyzed the effect of hM4Di activation on licking behavior and BG calcium between 1 and 2 hours after CNO administration. Upon intraperitoneal injection of CNO, capsaicin-induced licking behavior and BG flares were significantly reduced (Fig. 3d–f). In addition, we investigated the role of LC terminals in the cerebellar cortex in this behavioral response and BG flares. The total duration of capsaicin-induced licking behavior was reduced when C21 was locally applied through the cannula implanted in the cerebellar vermal lobule 4/5 of the LC-specific hM4Di-expressing mice (Fig. 3g). Moreover, capsaicin-induced BG flares were completely blocked (Fig. 3h, i). Taken together, these results demonstrate that capsaicin-induced acute tonic pain is associated with waves of BG flares as BG accumulate NA signals via LC activation. Moreover, we found that blocking these flares reduces the pain-related behavioral phenotype.

**Fig.3:**
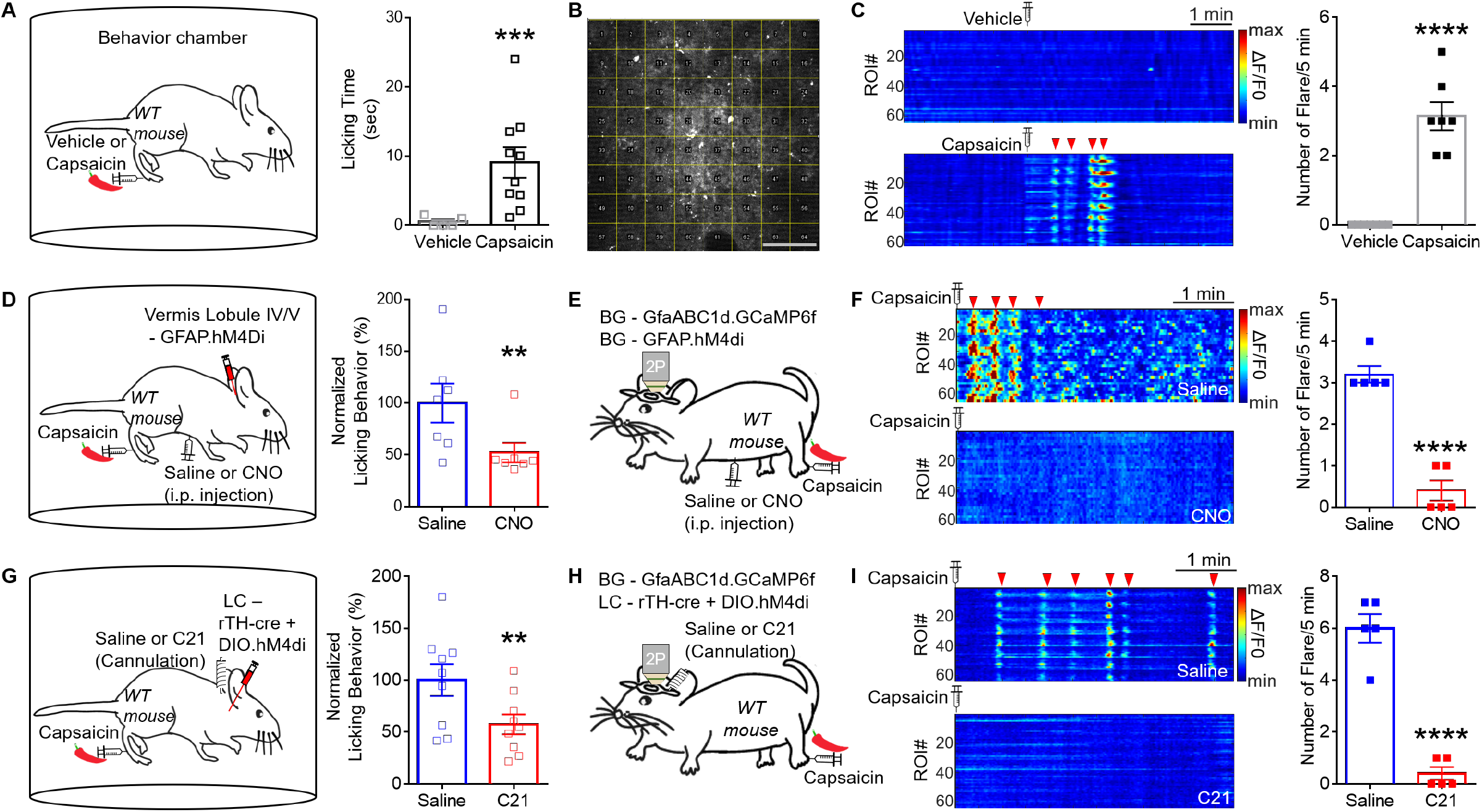
Capsaicin induces global BG calcium response (flare) associated with nocifensive behavior. **A,** Licking behavior in capsaicin-induced pain (n = 5 mice in vehicle group; n = 10 mice in capsaicin group; *** p = 0.0001; Two-way ANOVA). **B,** An example of ROI detection of BG using grid pattern. (8 by 8, 64 ROIs). **C,** Representative capsaicin-induced BG calcium response (above, vehicle; below, capsaicin; flare activity is indicated by red arrow head). BG calcium synchronous response as a “Flare” activity in capsaicin-induced pain (n = 7 mice in each group; **** p < 0.0001; Two-way ANOVA); Scale bar, 100 μm. **D,** Licking behavior in capsaicin-induced pain after BG hM4Di activation (n = 7 mice in each group; ** p = 0.0086; Two-way ANOVA). **E,** Schematic diagram showing *in vivo* two-photon imaging of BG calcium in response to hind-paw capsaicin injection in BG hM4Di expressing mouse. **F,** Representative capsaicin-induced BG calcium response (above, saline group; below, CNO group; flare activity indicated by red arrow). Flare activity in capsaicin-induced pain (n = 5 mice in each group; **** p < 0.0001; Two-way ANOVA). **G,** Licking behavior in capsaicin-induced pain after LC terminal hM4Di activation (n = 9 mice in each group; ** p = 0.0014; Two-way ANOVA). **H,** Schematic diagram showing *in vivo* two-photon imaging of BG calcium in response to hind-paw capsaicin injection in LC hM4Di expressing mouse. **I,** Representative of capsaicin-induced BG calcium response (above, saline group; below, C21 group; flare activity indicated by red arrow). Flare activity in capsaicin-induced pain (n = 5 mice in each group; **** p < 0.0001; Two-way ANOVA).

To manipulate BG flares based on the cell-type-specific molecular mechanisms, we investigated which type of NA receptors mediate capsaicin-induced BG flares and intraplantar capsaicin injection-evoked licking. We performed *in vivo* calcium imaging of BG as well as behavioral analysis with the local administration of NA receptor blockers (prazosin, an α1-AR blocker; yohimbine, an α2-AR blocker; propranolol, a β-AR blocker) through a cannula targeting the imaging area (Fig. 4a). Among the tested drugs, only prazosin completely blocked capsaicin-induced BG flares (Fig. 4b). Capsaicin-induced licking was also reduced in the prazosin-administration group (Fig. 4c). To evaluate whether this result arose from unintended effects of prazosin on the other cell types that express α1-AR, we measured the change in PC activity after prazosin application and found no effect (Fig. S4). The cerebellar slice recordings showed that BG calcium elevation during optogenetic activation of the LC terminal was completely blocked by application of prazosin through bathing solution (Fig. 4d–f). To further use cell-type-specific genetic manipulation, BG-specific α1-AR knockdown (KD) was achieved by injecting Cre-dependent α1 -AR shRNA virus into Glast-cre/ert2 mice, a transgenic strain specifically expressing Cre in BG (Fig. 4g, i) [32]. In the heterologous expression system, the calcium response of α1 -AR-KD glial cells was confirmed to be completely abolished in response to the α1-AR agonist but not the protease-activated receptor 1 agonist (Fig. S5a–c). Moreover, α1-AR protein expression was reduced by approximately 70% in the cerebellar cortex of the α1-AR KD group (Fig. S5d–f). Additionally, α1-AR KD completely blocked capsaicin-induced BG flares and significantly reduced capsaicin-induced licking compared to that in the sh-scramble-injected group (Fig. 4h, i). These results indicate that α1-AR on BG mediates the processing of pain information in the LC-cerebellum circuitry.

**Fig.4:**
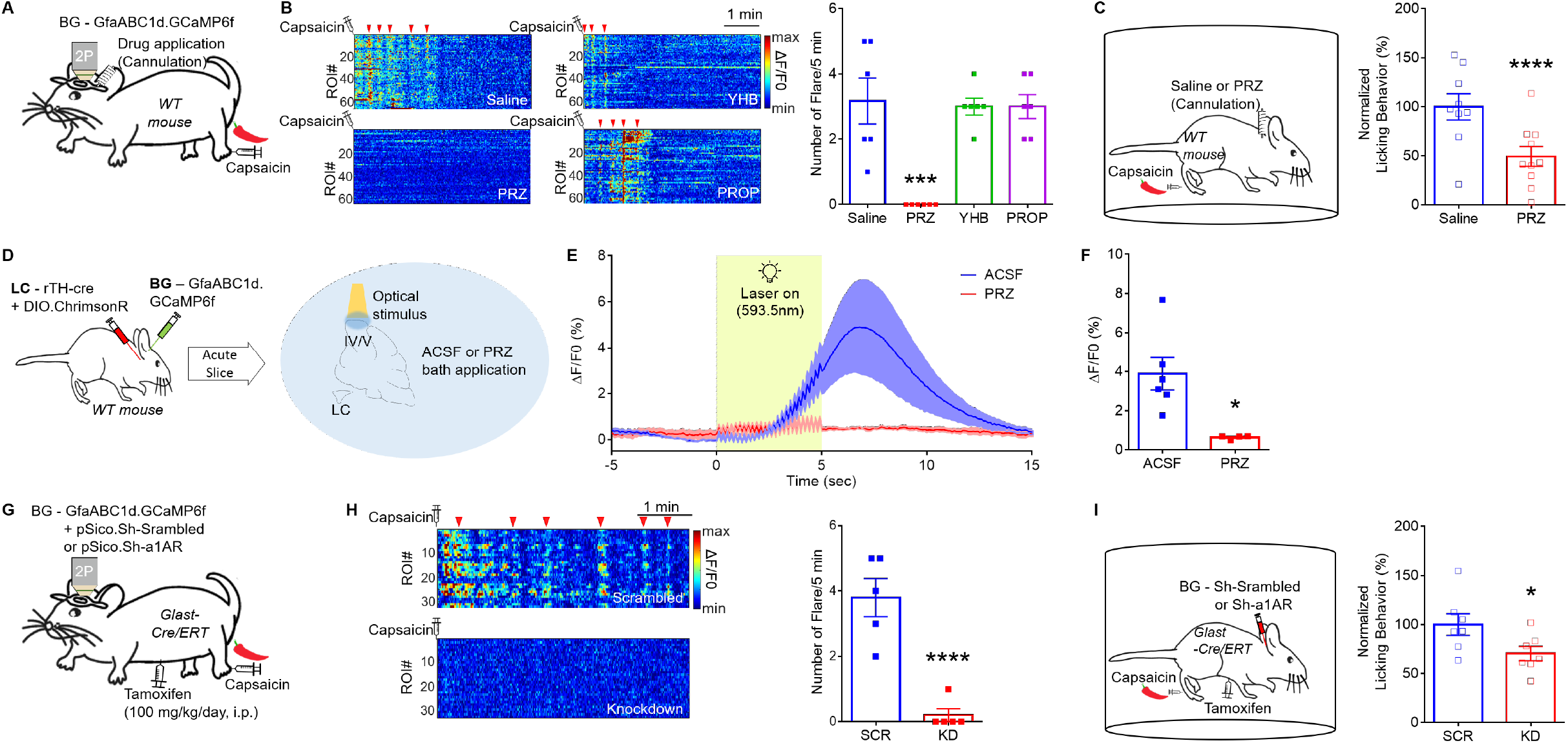
α1-AR mediates capsaicin-induced BG calcium flare and nocifensive behavior. **A,** Schematic diagram showing in vivo two-photon BG calcium imaging and drug application using the cannula simultaneously. **B,** Representative of capsaicin-induced BG calcium response (Saline, left above; Prazosin (PRZ), left below; Yohimbine (YHB), right above; Propranolol (PROP), right below; flare activity indicated by red arrowheads). Flare activity in capsaicin-induced pain (n = 6 mice in each group; *** p = 0.0003; Two-way ANOVA). **C,** Licking behavior in capsaicin-induced pain (n = 9 mice in saline group, n = 10 mice in prazosin group; **** p < 0.0001; Two-way ANOVA). **D,** Schematic showing slice imaging of BG calcium in response to LC terminal optogenetic modulation during prazosin bath application. **E,** Trace of BG calcium signal during LC optogenetic activation or control (ACSF, n = 6 slice in 5 mice; Prazosin, n = 4 slice in 2 mice; Line for Mean, shade for SEM; using 539.5 nm, 10 mW, 5 Hz, 5 sec). **F,** Decreased BG calcium activity by prazosin application (* p = 0.0146; Unpaired t-test). **G,** Schematic diagram showing *in vivo* two-photon imaging of BG calcium in response to capsaicin-injection in BG-specific genetic manipulation model mice. **H,** Representative of capsaicin-induced BG calcium response (above, scrambled group; below, knockdown group; flare activity indicated by red arrow head). Flare activity in capsaicin-induced pain (n = 5 mice in each group; **** p < 0.0001; Two-way ANOVA). **I,** Licking behavior in Capsaicin-induced pain (n = 7 mice in each group; * p = 0.0012; Two-way ANOVA).

We reasoned that BG flares affect pain-related licking behavior by modulating nearby neurons. In human brain imaging studies, noxious stimulations were found to activate the cerebellum [5–7]. The deep cerebellar nuclei (DCN), which are the output centers of the cerebellum, activate in response to noxious heat [5]. We hypothesized that BG activation inhibits nearby PCs in the cerebellar cortex, and in turn, PCs disinhibit DCN activity by decreasing their tonic inhibitory input to the DCN. To determine whether BG activation is responsible for the reduction in PC firing, we performed cell-attached *ex vivo* electrophysiology recordings with BG-specific chemogenetic manipulation in the cerebellar slices (Fig. 5a). The spontaneous firing rate of PCs was reduced when BG were activated using CNO application under hM3Dq expression. Inhibiting BG with hM4Di expression by CNO did not affect the spontaneous firing rate of PCs (Fig. 5b, c). *In vivo* electrophysiology recordings of cerebellar PCs indicated that capsaicin injection into the hind paw reduced the firing frequency. This reduction was blocked by microinjection of prazosin into the recoding site of the cerebellar cortex (Fig. 5d-f).

**Fig.5:**
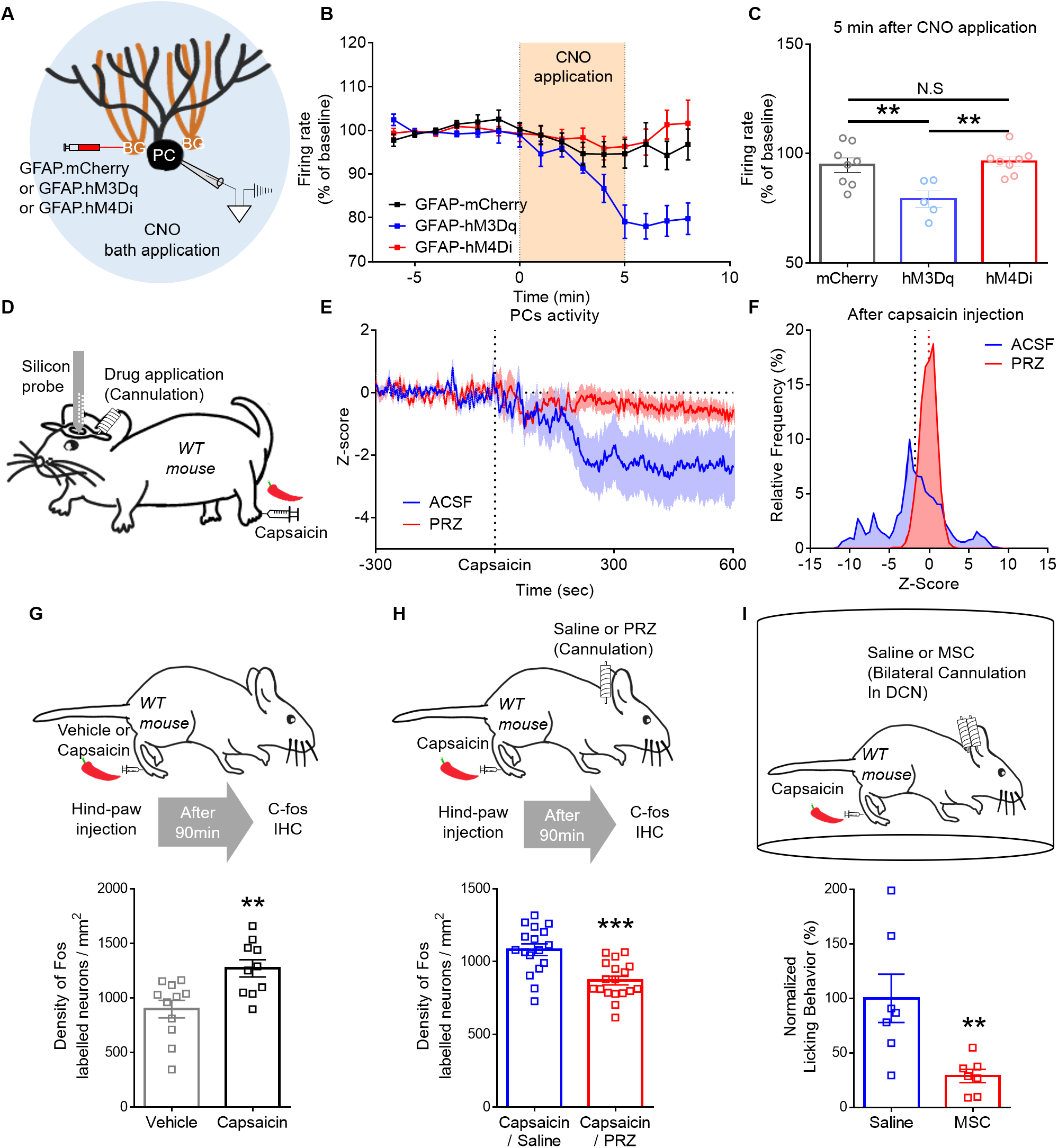
Manipulation of BG calcium regulates Purkinje cell activity and nocifensive behavior. **A,** Schematic diagram showing measurement of PC attached recording using BG chemogenetic manipulation. **B,** PC firing rate affected by BG activity modulated by CNO application (n = 8 slices in mCherry group; n = 5 slices in hM3Dq group; n = 8 slices in hM4Di group). **C,** PC firing rate after 5 min of CNO application (hM3Dq and mCherry, ** p = 0.0085; hM3Dq and hM4Di, ** p = 0.0039; hM4Di and mCherry, N.S. p = 0.9113; One-way ANOVA). **D,** Schematic diagram showing measurement of PC spontaneous firing using in vivo electrophysiology and drug application using cannula simultaneously. **E,** Representative of PC spontaneous firing in capsaicin-induced pain (Red, Prazosin application group; Black, ACSF application group). **F,** Changes in PC firing after capsaicin injection. **G,** c-fos expression data showing increase in DCN activity during capsaicin induced pain (n = 11 slices of 3 mice in vehicle group; n = 10 slices of 3 mice in capsaicin group; ** p = 0.0038; Unpaired t-test). **H,** c-fos expression data showing reduced in DCN activity during capsaicin induced pain by local prazosin injection to cerebellar lobule 4/5 (n = 17 slices of 3 mice in saline group; n = 18 slices of 3 mice in prazosin group; *** p = 0.0001; Unpaired t-test). **I,** Capsaicin-induced licking behavior after saline or muscimol injection in medial DCN (n = 7 mice for each group; *** p = 0.0003; Two-way ANOVA).

As prazosin reversed the capsaicin-induced reduction of PC firing, we speculated that the reduction of PC inhibitory input to the DCN during capsaicin-induced pain could be restored in the absence of BG involvement. Thus, the restored PC output may yield stronger inhibition of the DCN compared with that during the normal pain state. To confirm DCN activity during prazosin application, we examined the expression of the neuronal activity marker, c-Fos. We observed that c-Fos expression in the DCN increased during capsaicin-induced pain, and the prazosin group exhibited significantly lower c-Fos expression than the saline control group after capsaicin injection (Fig. 5g, h). Furthermore, to investigate the functional role of the DCN in pain processing, we applied muscimol, a GABA-A receptor agonist, to the DCN through the cannula, which inhibited capsaicin-induced licking (Fig. 5i). .00.

## Discussion

We found that BG in the cerebellar cortex play a central role in the computation of NA-driven noxious information. First, noxious information enters the cerebellum through the LC, and NA released from the LC terminals in the cerebellar cortex elevates BG calcium in an accumulative manner, demonstrating that BG encode the noxious information in the cerebellar cortex. Second, at the cellular level, the firing of PCs, which is the sole output of the cerebellar cortex, is modulated by BG calcium activity. Third, at the molecular level, the increased BG calcium activity and related regulation of PC activity in pain states are dependent on the α1-AR on BG. Finally, several interventions applied to the cerebellar neuron-glia network during intraplantar capsaicin injection reduced the expected nocifensive behavior. Through these findings, we have clearly elucidated the underlying mechanisms of nocifensive behavior on multiple levels and found that the cerebellum is actively involved in the computation of the pain state.

Previously, LC neurons were thought to be functionally homogeneous, broadly projecting noradrenergic fibers to the functionally diverse areas of the brain [33]. More recent studies suggest that LC neurons innervate anatomically and functionally distinct brain targets and can be grouped into several populations based on their preferred target [34]. In previous studies, the specific role and function of LC neurons was demonstrated by manipulating the LC terminal activity in the target brain areas [35–38]. The cerebellum is a target area of LC neurons, but its specific role was not previously understood. In this study, we dissected a novel role of LC terminals in the cerebellar cortex using two-photon microscopy and chemogenetics in a projection-specific manner. We found that calcium in the LC terminals in the cerebellar cortex increases in response to noxious stimulation (Fig. 1) and that nocifensive behavior can be modulated through chemogenetic inhibition of the LC terminal by locally applying C21 in the cerebellar cortex (Fig. 3g). This projection-specific approach demonstrates that the LC relays noxious information to the cerebellar cortex, contributing to nociception.

During cutaneous electrical stimulation, LC terminals in the cerebellar cortex responded with similar and repetitive amplitudes. In contrast, BG integrated NA signals and showed a gradual increase in calcium activity. The degree of BG calcium elevation was also affected by the frequency of stimulation (Fig. S2d), which could explain the temporal nature of astrocytes to control the state of neural networks in a distinct time range. Several reports have suggested that astrocytes can regulate neuronal activity and induce behavioral changes [22, 30, 39]. Likewise, temporal accumulation of noxious information in BG could enable animals to distinguish tonic pain from phasic pain and elicit coping responses to inevitable tonic pain. Furthermore, PC activity gradually decreased approximately 5 min after capsaicin injection or BG activation. This result suggests that PC activity could be progressively regulated during several flare events, indicating the presence of multiple temporal dynamics in the cerebellar circuit during noxious information processing.

We inhibited BG calcium activity using three crucial methods: (i) by activating exogenous BG-specific hM4Di to block overall BG activity (Fig. 3f, i); (ii) using prazosin, an α1-AR blocker, via a local pharmacological approach to confirm the molecular mechanism (Fig. 4b); and (iii) by inhibiting endogenous α1-AR via BG-specific α1-AR KD (Fig. 4g). Through these three methods, an increase in BG calcium under pain conditions was blocked and the accompanying nocifensive behavior was reduced. Previous studies have shown the relationship between BG calcium activity and motor behavior [14, 29, 45]; however, they did not evaluate the effect of suppressing BC calcium activity on motor behavior. In this study, we confirmed that neither BG-specific hM4Di nor BG-specific α1-AR KD had an effect on basal motor function (Fig. S6). Thus, we demonstrated that suppressing BG calcium activity does not change basal motor function but rather reduces nociceptive behavior, suggesting that BG calcium activity is crucial in the processing of noxious information.

We found that PC firing was suppressed after capsaicin injection via the NA-BG pathway. Although we revealed the BG-PC relationship using cell-type-specific chemogenetic manipulation (Fig. 5d-f), the underlying mechanism of the interaction between BG and PCs remains unclear. Nevertheless, several previous studies have attempted to explain how BG regulate PC activity [19–21, 41, 42]. First, BG can influence PC activity through the release of gliotransmitters. The release of glutamate from BG activates the molecular layer interneurons via the activation of the NMDA receptor [41]. GABA release from BG through the Bestrophin-1 channel can also directly inhibit PCs [42], and tonic inhibitory currents can hyperpolarize PCs. Second, through glial regulation of extracellular ionic composition [20, 43–45], BG activation can lower the extracellular potassium concentration via potassium uptake. As a result, the membrane potential may become more negative, suppressing the firing of PCs. These two mechanisms may underlie the BG-dependent reduction of PC activity in the pain state.

We further evaluated how BG activation mediates behavioral responses to noxious stimuli. We hypothesized that suppressed PC activity during capsaicin-induced pain may disinhibit DCN activity [46, 47]. Using c-fos staining, we demonstrated that DCN activity increased after intraplantar capsaicin injection and was restored by the blockade of α1 -AR. Activated (disinhibited) DCN may deliver noxious information, which has been computed in the cerebellar microcircuit, to DCN-connected brain regions involved in nocifensive behavior, such as the periaqueductal gray, parabrachial nucleus, and zona incerta [48–51]. We also investigated the engagement of DCN in nocifensive behavior and found that muscimol-induced DCN inhibition significantly reduced nocifensive behavior. In summary, we examined the complex cerebellar circuit, including the neuro-modulatory system and neuro-glial interactions involved in pain information processing, from NA-BG to the DCN.

Clinically, NA inhibitors have been tested in patients with complex regional pain syndrome [52, 53]. However, explaining the exact mechanism of these analgesic effects is difficult owing to the limitations of systemic application and side effects. In this study, blocking α1-AR specifically in the cerebellum resulted in a significant analgesic effect, implicating an noble approach to the cerebellum with the α1 -AR blockade. Notably, capsaicin-induced licking behavior was reduced by local manipulation of the cerebellum, without affecting motor function (Fig. S7). Our study clearly showed that pain could be modulated by blocking only BG in the cerebellar vermal lobule 4/5 without the risk of adverse or systemic effects. Eventually, our research will provide a basis for a new approach to controlling pain by applying optogenetics, chemogenetics, and electroceutical tools to specific cells in specific areas.

## Acknowledgements

We thank Geehoon Chung, Dong Cheol Jang, Kyoung-Doo Hwang and Hyun Geun Shim for critical discussions on the experiments and writing. We thank Dr. Magdalena Götz (Helmholz Center Munich, Institute Stem Cell Research, Munich, Germany) and Professor K. Tanaka (Tokyo Medical and Dental University, Tokyo, Japan) for providing GLAST-CreERT2 transgenic mice

## Author contributions

Conceptualization: SHK, SR, SKK, SJK

Investigation: SHK, JL, SR, SbK, MJ, JHL, JYH

Funding acquisition: SKK, SJK

Supervision: YL, ES, CJL, SK, SKK, SJK

Writing – original draft: SHK, JL, SKK, SJK

Writing – review & editing: SHK, JL, SR, JS, SKK, SJK

## Data and materials availability

All software or algorithm used in this study is available and listed in the Key Resources Table. The All data and codes associated with this study are presented in the paper, supplementary materials. The data that support the findings of this study are available upon reasonable request from the lead contact

## Supplementary Materials

Materials and Methods

Figs. S1 to S9

References

## Supplementary Materials

### Materials and Methods

#### Key resources table

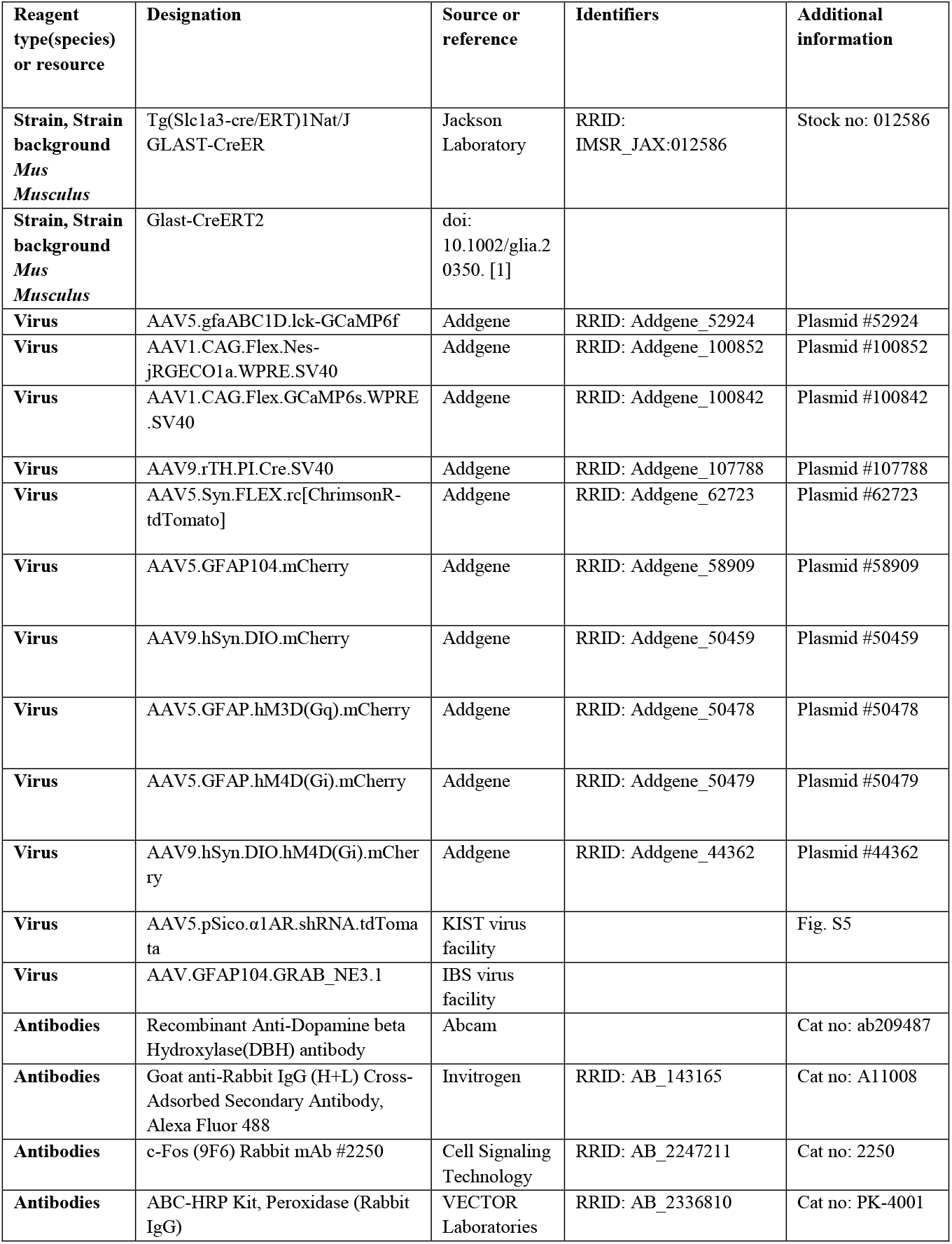
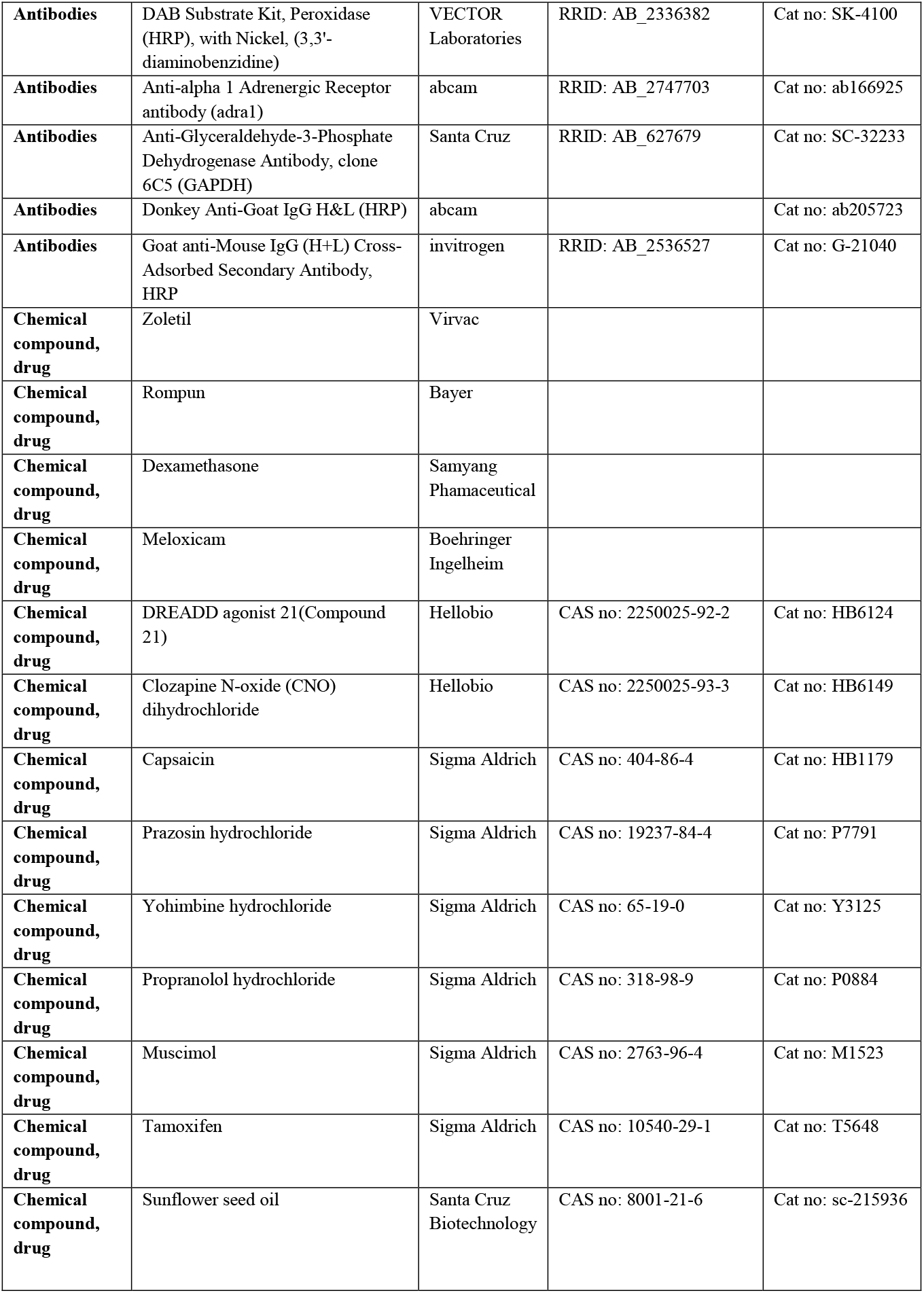
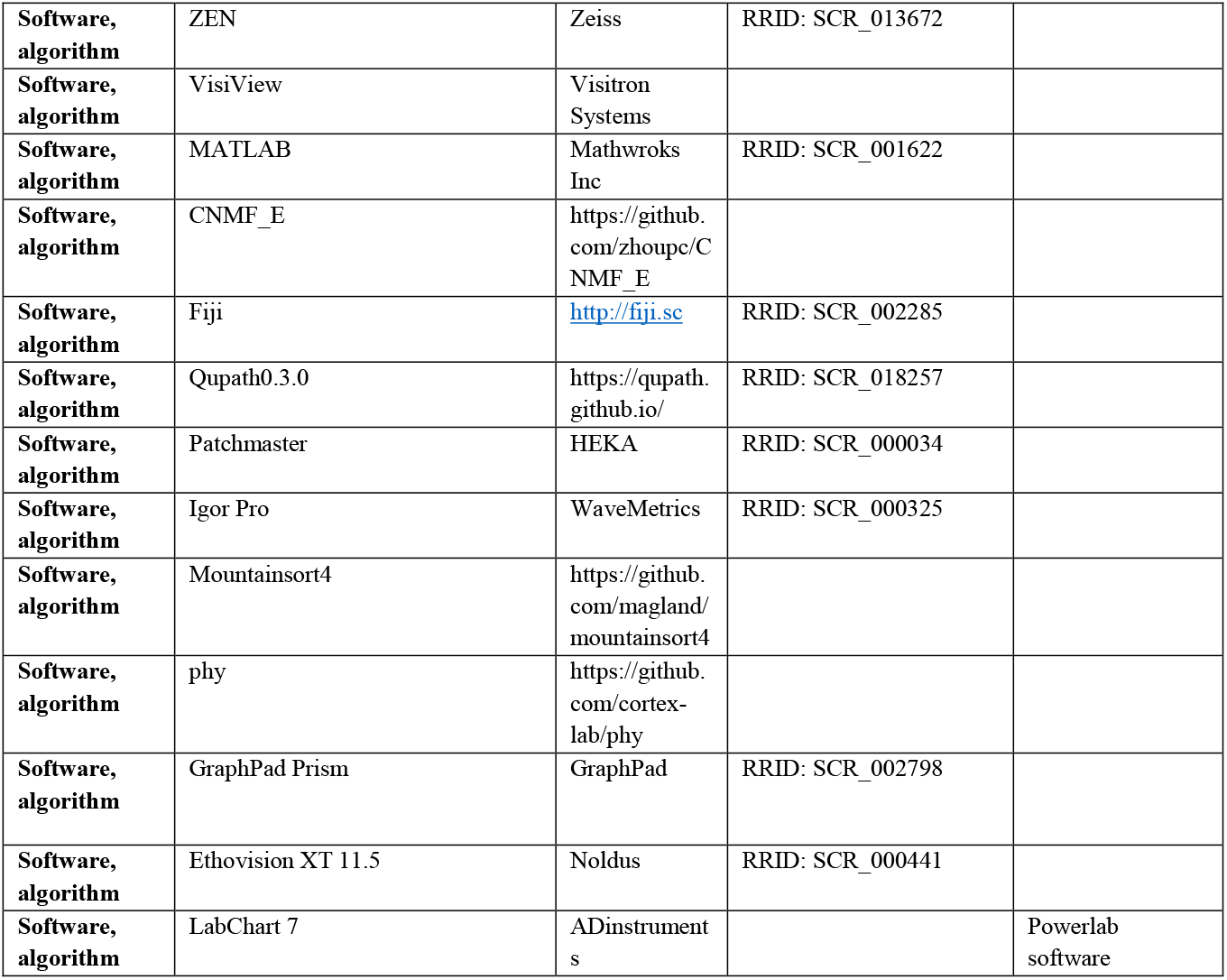

#### Animals, craniotomy

The experimental processes were approved by the Seoul National University Institutional Animal Care and Use Committee and performed under the guidelines of the National Institutes of Health and the Animal Care Committee of Yamanashi University (Chuo, Yamanashi, Japan). Seven- to ten-week-old wild-type or astrocyte-specific Tamoxifen inducible cre expressing mouse line, TG(Slca3-cre/ERT)1Nat/J (Jackson Laboratory, ME, USA) as known as Glast-CreER. Glast-CreERT2 mice were provided by Dr. Magdalena Götz (Helmholz Center Munich, Institute of Stem Cell Research, Munich, Germany) and by Dr. K. Tanaka (Tokyo Medical and Dental University, Tokyo, Japan) [1]. mice were anesthetized with intraperitoneal injections of Zoletil/Rompun mixture (30 mg / 10 mg/kg) or under 1.5-2 % isoflurane. A small craniotomy was made over lobule 4/5 of the cerebellar vermis/paravermis according to previous descriptions but with some modifications [2, 3]. In short, after placing the anesthetized mouse on a stereotaxic frame (Narishige, Tokyo, Japan), the skin was incised, and bone was removed with a no.11 surgical blade. To minimize edema and related inflammation, dexamethasone (0.2 mg/kg) and meloxicam (20 mg/kg) were administered by subcutaneous injection. A metal ring for head fixation was attached with Superbond dental cement (Sun Medical, Japan). Then, a 1.4 × 2.3 mm size glass coverslip (Matsunami, Japan) was tightly placed on the cortex and fixed by applying cyanoacrylate glue (Vetbond, 3M), as previously described [3].

#### Viral Expression and injection

For BG-specific GECI expression, 100-200 nl virus solution of 3-5 × 10^10^ genome copies containing AAV5.gfaABC1d.Lck-GCaMP6f (Addgene, MA, USA) were injected at two or three sites in the craniotomy site at the cerebellar cortex (Vermis Lobule 4/5) of the WT mice with a beveled glass pipette (5 MΩ). For GECI expression in LC, the virus was injected into the bilateral LC by making a small hole in the skull (posteriorly 5.4 mm and bilaterally 1.1 mm from bregma) with a dental drill. The glass pipette was set vertical angle for the skull. After approaching a 3.5 mm depth, virus solution containing 100-200 nl mixture of AAV9.rTH.PI.Cre.SV40 and AAV1.CAG.GCaMP6s.WPRE.SV40 or AAV1.CAG.Flex.Nes-jRGECO1a.WPRE.SV40 (Addgene, MA, USA) was injected with a Picopump at 5 nl/sec. The pipettes were left in place for 10 min before they were removed to minimize backflowing. For red-shift GECI expression in BG, AAV1.CAG.Flex.Nes-jRGECO1a.WPRE.SV40 was injected into Glast-CreER (Jackson Laboratory, ME, USA) mice. For NE sensor expression in BG, AAV.GFAP104.GRAB_NE3.1 (IBS virus facility, Daejeon, Korea) was injected into wild-type mice. Grab_NE working mechanism has been described previously [4]. For Chemogenetic expression in BG, AAV5.GFAP.hM3D(Gq).mCherry or AAV5.GFAP.hM4D(Gi).mCherry was injected into wild-type mice. For mCherry expression in BG, AAV5.GFAP104.mCherry was injected into wild-type mice. For Chemogenetic expression in LC, a mixture of AAV9.rTH.PI.Cre.SV and AAV9.hSyn.DIO.hM4D(Gi).mCherry was injected into wild-type mice. For Optogenetic expression in LC, a mixture of AAV9.rTH.PI.Cre.SV and AAV5.Syn.FLEX.rc[ChrimosonR-tdTomato] was injected into wild-type mice. For mCherry expression in LC, a mixture of AAV9.rTH.PI.Cre.SV40 and AAV9.hSyn.DIO.mCherry was injected into wild-type mice. For α1-AR knockdown expression in BG, AAV5.pSico. α1AR.shRNA.tdTomato (KIST virus facility, Seoul, Korea) was was injected into cre-activated Glast-CreER or Glast-CreERT2 mice by repeating tamoxifen (100 mg/kg) for 5 days.

#### Two-photon microscopy and chronic window imaging

In vivo imaging experiments of genetic manipulation model using Glast-CreERT2 were performed using an Olympus Fluoview FVMPE-RS two-photon laser scanning microscope (Olympus, Tokyo, Japan) equipped with Maitai HP DS-OL (Spectra-Physics, Santa Clara, CA) and InSight DS-OL (Spectra-Physics). We used the laser at 920 nm and a 495-540 nm bandpass pass emission filter for GCaMP6f and the laser at 1100 nm and a 575-645 bandpass emission filter for tdTomato. Images were gathered with a 25 × water immersion lens (XPLN25XWMP2) with a numerical aperture of 1.05 (Olympus). All other experiments were performed with confocal microscopy was performed with a laser scanning multiphoton microscope (Zeiss LSM 7 MP, Carl Zeiss, Jena, Germany) equipped with non-descanned detectors (NDD). Excitation was carried out with a mode-locked titanium:sapphire laser system (Chameleon, Coherent, Santa Clara, CA, USA) operating at a 900 nm wavelength for GCaMP6 and GRAB_NE using GFP filter and 980 nm for jRGECO1a and mCherry using RFP filter. Generally, objective W Plan-Apochromat 20×, 1.0 numerical aperture (Carl Zeiss) was used. Images were acquired using ZEN software (Zeiss Efficient Navigation, Carl Zeiss) and processed using a custom-written MATLAB (MathWorks) script. High-resolution of 512 × 512-pixel reference images were acquired at a rate of 8 s per frame in the molecular layer (around 20–60 μm from the dura). For BG Ca^2+^ imaging, 2-5 Hz speed time-lapse scanning was performed with a 512 × 512 resolution window at 50-100 μm from the dura. For LC terminal Ca^2+^ imaging, 5-8 Hz speed time-lapse scanning was performed with a 512 × 256 or 512 × 512 resolution window at 50-100 μm from the dura. For chronic window imaging, 14-21 days after chronic window surgery, the mice were imaged under 1-1.5 % isoflurane anesthesia with head-fixed using a clamp and custom-made metal rings.

#### BG and LC ROI detection and calcium analysis

Since it is difficult to specify each BG process, an ROI was constructed with 64 grids for 512-512-pixel images and 32 grids for 512-256-pixel images and analyzed [5]. CNMF_E was used to select ROIs for the LC terminal in the cerebellar vermis lobule 4/5 [6] and coordinate ROI values obtained through CNMF_E were confirmed through image j, and ROIs in the same location were manually combined (Fig. S9). Customized scripts in MATLAB were used to analyze the calcium transient signals. Calcium signal amplitudes were calculated as ΔF/F0 (ΔF = F - F0) for each cell. F0 means the baseline fluorescence signal calculated by averaging the lowest 10 % of all fluorescence signals from individual traces. The change value of the maximum value of calcium signal from noxious electrical stimulation was confirmed by comparing it with the baseline. To characterize the flare, BG calcium activation was confirmed through noxious electrical stimuli before capsaicin-induced pain experiments. After confirming the calcium response, the period of at least 60 % synchronization and calcium activation as much as the amount of calcium from electrical stimulation of 5-10 seconds or more is specified as a flare.

#### Noxious Stimulation

To deliver noxious electrical stimulation, a needle with a blunt tip is inserted into the plantar side of the hind paw, and stimulation with 3 Hz frequency, 0.2 ms duration, and 0, 1, 10 mA power is provided for 10 seconds using Powerlab 4/25T (ADInstruments, New South Wales, Australia) device.

For actual pain stimulation, 0.01% capsaicin solution dissolved in 20 % ethanol (in PBS) was used. Licking behavior was observed when 10 μl capsaicin solution was injected into the hind paws, which disappeared within 10 minutes each, respectively.

#### Drug Application

A unilateral cannula was installed to inject the drug only into the target site, cerebellar vermis lobule 4/5 (−6.0 mm anteroposterior from bregma, midline). Drugs such as AR blocker (Prazosin, 100 μM; yohimbine, 100 μM; propranolol, 100 μM) or C21 (1 mM) used in the experiment were injected through this cannula with a syringe pump (Pump 11 Elite Infusion Only; Harvard apparatus), and 0.5 μl was slowly injected for about 10 minutes.

A bilateral cannula was installed to inject the drug only into the target site, deep cerebellar nuclei (−6.2 mm anteroposterior from bregma, ±1 mm mediolateral from bregma, −3.2 mm dorsoventral from the surface of skull). Drugs such as GABA A agonist (Muscimol, 100 μM) used in the experiment were injected through this cannula with a syringe pump (Pump 11 Elite Infusion Only; Harvard apparatus), and 0.5 μl was slowly injected for about 10 minutes.

#### Chemogenetic modulation

For BG chemogenetic modulation, BG-specific virus AAV5.GFAP.hM4D(Gi).mCherry, was injected into cerebellar vermis lobule 4/5. To prevent systemic activation by intraperitoneal injection of CNO (5 mg/kg), the virus was locally injected into the cerebellar vermis lobule 4/5 and acted only on the expressed lobules. Behavioral experiments and calcium imaging were performed 1hr after CNO intraperitoneal injection by validation of hM4Di activation (Fig. S3).

For LC chemogenetic modulation, LC-specific virus, mixture AAV9.rTH.PI.CRE.SV40 and AAV9.hSyn.hM4D(Gi).mCherry, was injected into LC. To prevent systemic activation by C21, a cannula was installed in the cerebellar vermis lobule 4/5, and C21 (1 mM, 0.5 μl) was locally infused through the cannula. Then, the virus is expressed in the whole brain, but the effect of C21 appears only locally in the cerebellum. Behavioral experiments and calcium imaging were performed 10 min after C21 infusion.

To determine the effect on PC firing through BG chemogenetic modulation, we measured spontaneous PC firing rate from cerebellar slices with CNO application. CNO (10 μM) was perfused for 5 min following a 5 min baseline period.

For BG-specific Gq activation of cerebellar vermis lobule 4/5, AAV5-GFAP.hM3D(Gq).mCherry, was injected. In the same way as BG hM4Di activation, CNO was administered intraperitoneally, and behavioral experiments were performed 1 hour later (Fig. S7).

For BG-specific Gq activation of other regions in the cerebellum, we selected crus 1/2 in the cerebrocerebellar region and cerebellar vermis lobule 9/10 in the vestibulocerebellar region and injected the virus into each of these regions. As with the above BG chemogenetic modulation, the behavioral experiment was performed 1 hour after CNO intraperitoneal administration (Fig. S8).

#### Behavioral test

##### Capsaicin-induced pain test

To measure capsaicin-induced licking behavior, mice were placed in a cylindrical chamber with only the bottom transparent, and the bottom surface was acquired with a camera. After putting a mouse in the chamber and measuring the baseline for 10 minutes, the mouse was taken out and capsaicin is quickly injected into the hind paw, then put back into the chamber and taken for an additional 20 minutes. The first 10 minutes of the 20 minutes indicate the capsaicin phase, and after the 10 minutes indicate the recovery phase. We watched the video we recorded and manually recorded how many licks the injected paw was.

##### Open Field Test

Place the mouse in a chamber 40 cm x 40 cm, acquired video for 20 minutes, and measure the total distance traveled between various measurable indicators. In this case, the mouse should be the first time to experience the chamber. Mice behavior was recorded using a mouse-tracking software (EthoVision XT 11.5, Noldus, Netherlands)

##### Gait test, and Ledge test

This method was used with a slight modification of the previously described method [7]. The behavioral patterns of mice were scored on a scale of 0-3, with 3 as normal to 0 as abnormal. First, for measuring the Gait, place the mouse on a flat surface with its head facing away from the investigator and observe the mouse from behind as it walks. Scoring is done according to the following criteria. Score 3 – the mouse moves normally, with its body weight supported on all limbs, with its abdomen not touching the ground, and with both hindlimbs participating evenly, Score 2 - it shows a tremor or appears to limp while walking, Score 1 - it shows a severe tremor, severe limp, lowered pelvis, or the feet point away from the body during locomotion (“duck feet”), Score 0 – the mouse has difficulty moving forward and drags its abdomen along the ground. Next, for measuring the Ledge test, Place the mouse on the cage’s ledge and observe the mouse as it walks along the cage ledge and lowers itself into its cage. Scoring is done according to the following criteria. Score 3 – walk along the ledge without losing its balance, and will lower itself back into the cage gracefully, using its paws, Score 2 – the mouse loses its footing while walking along the ledge, but otherwise appears coordinated, Score 1 – it does not effectively use its hind legs, or lands on its head rather than its paws when descending into the cage, Score 0 – it falls off the ledge, or nearly so, while walking or attempting to lower itself, or shakes and refuses to move at all despite encouragement

Behavioral indicators were normalized according to capsaicin and each control group. All experiments were performed in a blinded manner, comparing the control group with the experimental group, respectively (Fig. S6).

#### Slice electrophysiology, optogenetic modulation, and imaging

An acute brain slice preparation and an electrophysiological experiment were carried out as previously described [8]. 5 to 9-week-old mice were anesthetized by isoflurane and briefly decapitated. Then, 250 μm-thick sagittal slices of the cerebellar vermis were obtained from mice using a vibratome (VT1200S, Leica). The ice-cold cutting solution contained 75 mM sucrose, 75 mM NaCl, 2.5 mM KCl, 7 mM MgCl_2_, 0.5 mM CaCl_2_, 1.25 mM NaH_2_PO_4_, 26 mM NaHCO_3_, and 25 mM glucose with bubbled 95% O_2_ and 5% CO_2_. The slices were immediately moved to artificial cerebrospinal fluid (ACSF) containing 125 mM NaCl, 2.5 mM KCl, 1 mM MgCl_2_, 2 mM CaCl_2_, 1.25 mM NaH_2_PO_4_, 26 mM NaHCO_3_, and 10 mM glucose with bubbled 95 % O_2_ and 5 % CO_2_. Then, they were recovered at 32 °C for 30 min and at room temperature for 1 hr. All recordings were performed within 8 hr after recovery.

The brain slices were placed in a submerged chamber with perfusion of ACSF for at least 10 min before recording. Cell-attached recordings were made at 29.5-30 °C. We used recording pipettes (3–4 MΩ) filled with (in mM): 9 KCl, 10 KOH, 120 K-gluconate, 3.48 MgCl_2_, 10 HEPES, 4 NaCl, 4 Na_2_ATP, 0.4 Na_3_GTP, and 17.5 sucrose (pH 7.25). Electrophysiological data were acquired using an EPC9 patchclamp amplifier (HEKA Elektronik) and PatchMaster software (HEKA Elektronik) with a sampling frequency of 20 kHz, and the signals were filtered at 2 kHz. All electrophysiological recordings were acquired in lobule 4/5 of the cerebellar central vermis. The spontaneous firing rate was analyzed using Igor Pro (WaveMetrics) and normalized by the average firing rate during the baseline period.

For BG Ca^2+^ or NE sensor imaging with LC optogenetic modulation and validation, slices were prepared from mice that had AAV5.gfaABC1d.Lck-GCaMP6f or AAV.GFAP104.GRAB_NE3.1 injected into their cerebellar vermis lobule 4/5, and AAV9.rTH.PI.Cre.SV40 and AAV5.Syn.FLEX.rc[ChrimsonR-tdTomato] was injected into their LC 3-4 weeks before the preparation. Using OptoPatcher (A-M systems) for slice recording with optogenetic manipulation, a 593.5 nm laser was used with 0-10 mW power, 1-5 Hz frequency, 50ms pulse width, 5 sec duration and then the signal from BG was acquired at 5 Hz with a scientific CMOS camera (Prime, Photometrics). All experiments were performed in duplicate or triplicate and traces of each manually selected cell were analyzed. Fluorescence images were acquired with VisiView software (Visitron Systems GmbH, Germany). The ROIs were manually detected in a field of view (Molecular layers within the image), and signals were extracted and converted to a normalized value (dF/F0). The value of F0 was calculated using the median value of the 5 and 15 percentile fluorescence in a sliding time window by each ROI.

#### In Vivo Electrophysiology

In vivo recordings were performed using anesthetized, head-fixed mice. Cranial window (1 × 3 mm) was made at least 1 week after cannula and ring implant under 1-1.5 % isoflurane. After head fixation, a custom-made 32-channel silicon probe was gradually inserted into vermis lobule 4/5 using a single-axis manipulator (SOLO, Sutter instrument). After confirming PC activities by complex spikes, ACSF or CNO was injected through a cannula. After at least 30 min of stable recording, capsaicin was injected at the hind paw ipsilateral side of the recording site. Single units were isolated using Mountainsort4 and manually curated using phy. Two types of band-pass filters were used for detecting simple spikes and complex spikes, 300-6000 Hz and 10-200 Hz, respectively. The firing rate was normalized to baseline activity before capsaicin injection using a z-score transformation. Increase or decrease units were defined as an average z-score after capsaicin injection > 3 or < −3.

#### Immunohistochemistry

Anesthetized LC specific mCherry expressing mice were perfused with PBS and again with 4 % paraformaldehyde (PFA; T&I). Brains were taken out and fixed in 4 % PFA overnight. After embedding the Frozen Section Compound FSC; LeicaBiosystems), we obtained 30-μm-thick sagittal slices on slides by using Cryotome (HM525 NX; Thermo Fisher Scientific). FSC was removed with PBS. Afterward, slices were permeabilization with a PBS-T (0.3 % Triton X-100) for 3 repetitions of 10 min at room temperature. Next, slices were blocked with a serum solution containing PBS-T and 5 % Bovine Serum Albumin BSA; Sigma Aldrich) for 3 repetitions of 1hr at room temperature. The slices were then incubated overnight at 4 °C with diluted primary antibody, anti-Dopamine beta-hydroxylase (anti-Rabbit, 1:500; Abcam). After washing in PBS-T, a fluorescence-conjugated secondary antibody, Alexa Fluor-488 (anti-rabbit, 1:500; Invitrogen), was used to treat the slices for 2 hr at room temperature. Primary antibodies and secondary antibodies were diluted in a blocking solution, respectively. Images were acquired and processed using a confocal microscope (Zeiss LSM 7 MP, Carl Zeiss, Jena, Germany) and Zen software (Zeiss).

#### Estimation of c-fos expression

To measure the c-fos expression in the DCN, DAB staining was performed following the manufacturer’s instructions (Vector Laboratories). It is mostly similar to the above-described immunohistochemistry, but there are some differences. First, Mice used for capsaicin-induced pain and prazosin cerebellar application experiments were perfused with PBS after 90 minutes of capsaicin injection with maximally increased c-fos. Second, the slices were then incubated for 48 hr at 4 °C with diluted primary antibody for c-fos (9F6) (anti-rabbit, 1:1000; Cell Signaling). Third, Signals were developed with the Vectastain ABC kit (PK-4001, Vector Laboratories), and DAB reagents (SK-4100, Vector Laboratories). Last, after washing in PBS and blocking with BSA, a biotinylated secondary antibody (anti-rabbit, 1:200; Vector Laboratories), was used to treat the slices for 24 hr at room temperature. Images were acquired and processed using a fluorescence microscope (IX53, Olympus), and scanned images were analyzed automatically using QuPath [9]. The measured image was 400 pixels (px) per 100 μm. The detection of nuclei was performed with the following settings: Requested pixel size Background radius 5 px, Median filter radius 3 px, Sigma 1.5 px, Minimum area 1 px^2^, Maximum area 400 px^2^, Intensity threshold 0.1, max background intensity 1.5-2, cell expansion 2 px split by shape. The numbers of cells and fluorescent spots were calculated through QuPath scripts. Estimation of the c-fos expression in the DCN was performed by counting the c-fos-positive cells per unit area (mm^2^).

#### Western blotting

Cerebellar vermis lobule 4/5 was dissected in a BG-specific α1-AR KD model mouse (Fig. S5d-e). Each cerebellar vermis lobule 4/5 was homogenized in 75 μl lysis buffer containing 10 mM Tris-HCl (pH 6.8) buffer, 1.6 % SDS, protease inhibitor, and phosphatase inhibitors. Samples were run on a 12 % bis tris gel and transferred to a PVDF membrane for other proteins. After blocking in 3 % BSA in 0.1 % TBST, membranes were probed with primary antibody (goat anti-ADRA1, 1:2000, Abcam, ab166925; mouse anti-GAPDH, 1:1,000, Santa Cruz Biotechnology, sc32233) overnight at 4 °C. After washing 3 times in 0.1 % TBST, membranes were probed with horseradish peroxidase-conjugated secondary IgG for 1 h at room temperature. Signals from membranes were detected by using an ECL chemiluminescence substrate kit (Thermo Pierce). Proteins were normalized to GAPDH, and phosphorylated proteins were normalized to their respective total proteins.

#### Statistics

Graph plotting and statistical analysis were carried out with GraphPad Prism (GraphPad Software Inc, CA). The hypothesis was tested by two-way ANOVA followed by post hoc Tukey’s test, for multiple comparisons using either GraphPad Prism or MATLAB. Paired and unpaired t-tests between sample pairs were carried out. Results were considered significant if the P-value was below 0.05. Asterisks denoted in the graph indicate the statistical significance. * means p-value < 0.05, ** < 0.01, *** < 0.001, and **** < 0.0001. The test name and statistical values are presented in each figure legend.

**Fig. S1.**
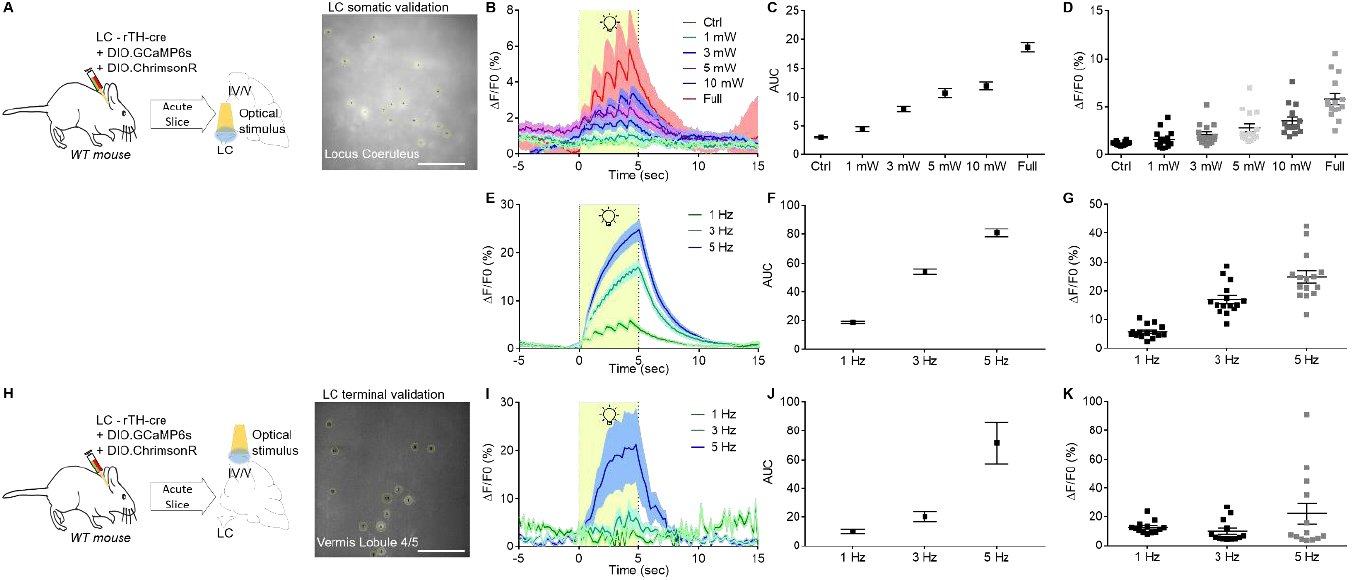
Validation of optogenetic modulation in LC neuron-specific manner. A, Schematic diagram showing virus injection for validation of LC somatic optogenetic modulation. (n = 15 ROIs). B, Validation of laser stimulation (593.5nm, 5 Hz, 5 sec) strength through LC soma activation and calcium response. C, AUC according to laser stimulation power. D, Peak of LC somatic calcium response according to laser stimulation power. E, Validation of laser stimulation frequency (593.5nm, 10 mW, 5 sec) through LC soma activation and calcium response. F, AUC according to laser stimulation frequency. G, Peak of LC somatic calcium response according to laser stimulation frequency. H, Schematic diagram showing virus injection for validation of LC terminal optogenetic modulation. (n = 13 ROIs). I, Validation of laser stimulation frequency (593.5nm, 10 mW, 5 sec) through LC terminal activation and calcium response. J, AUC according to laser stimulation frequency. K, Peak of LC terminal calcium response according to laser stimulation frequency.

**Fig. S2.**
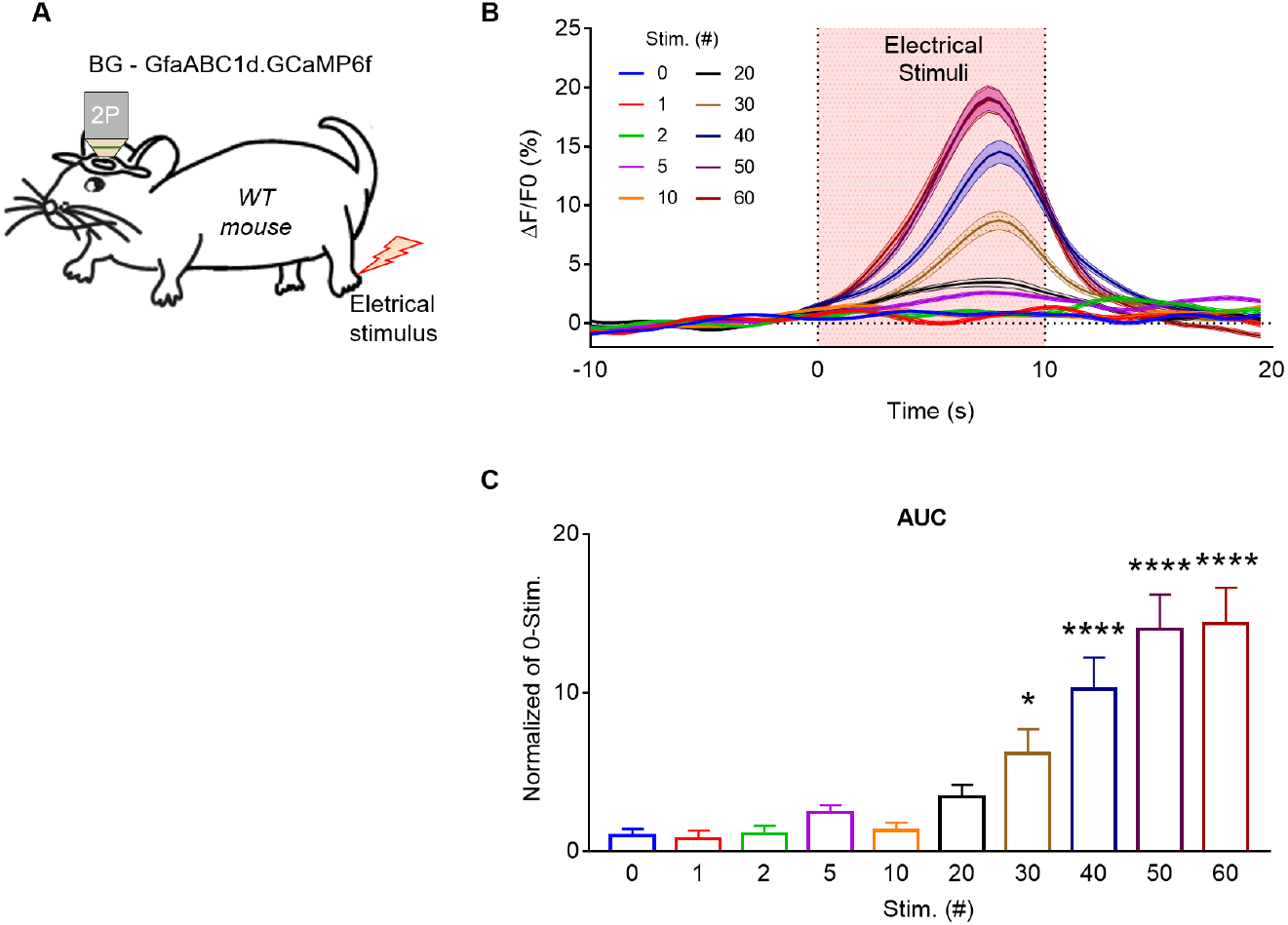
BG accumulates NE signals to produce calcium signals. A, Schematic diagram showing *in vivo* two-photon BG calcium imaging using hind-paw electrical stimulation. B, BG calcium response according to the number of stimulations (n = 192 grids in 6 mice). C, AUC of BG calcium response according to the number of stimulations (0 and 1 – 20 stim., N.S.; 0 and 30 stim., * p=0.0433; 0 and 40-60 stim., **** p=0.0001; One-way ANOVA).

**Fig. S3.**
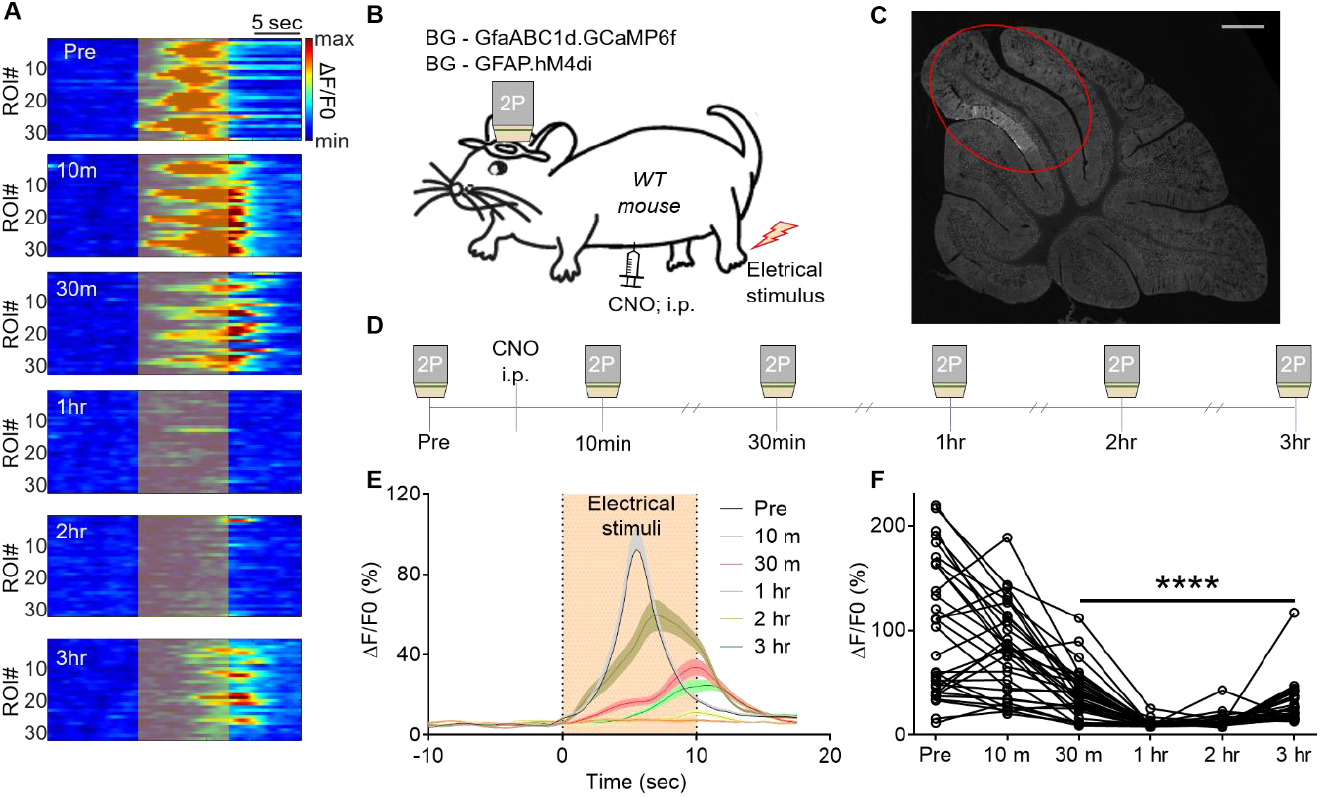
Validation of BG hM4Di activation *in vivo*. A, Representative BG calcium response before and after (~3 hr) CNO i.p. injection during hind-paw electrical stimuli. B, Schematic diagram showing *in vivo* two-photon BG calcium imaging using hind-paw electrical stimulation in BG hM4Di expressing mouse. C, Virus expression in cerebellar vermis lobule 4/5. D, Imaging schedule before and after CNO injection.

**Fig. S4.**
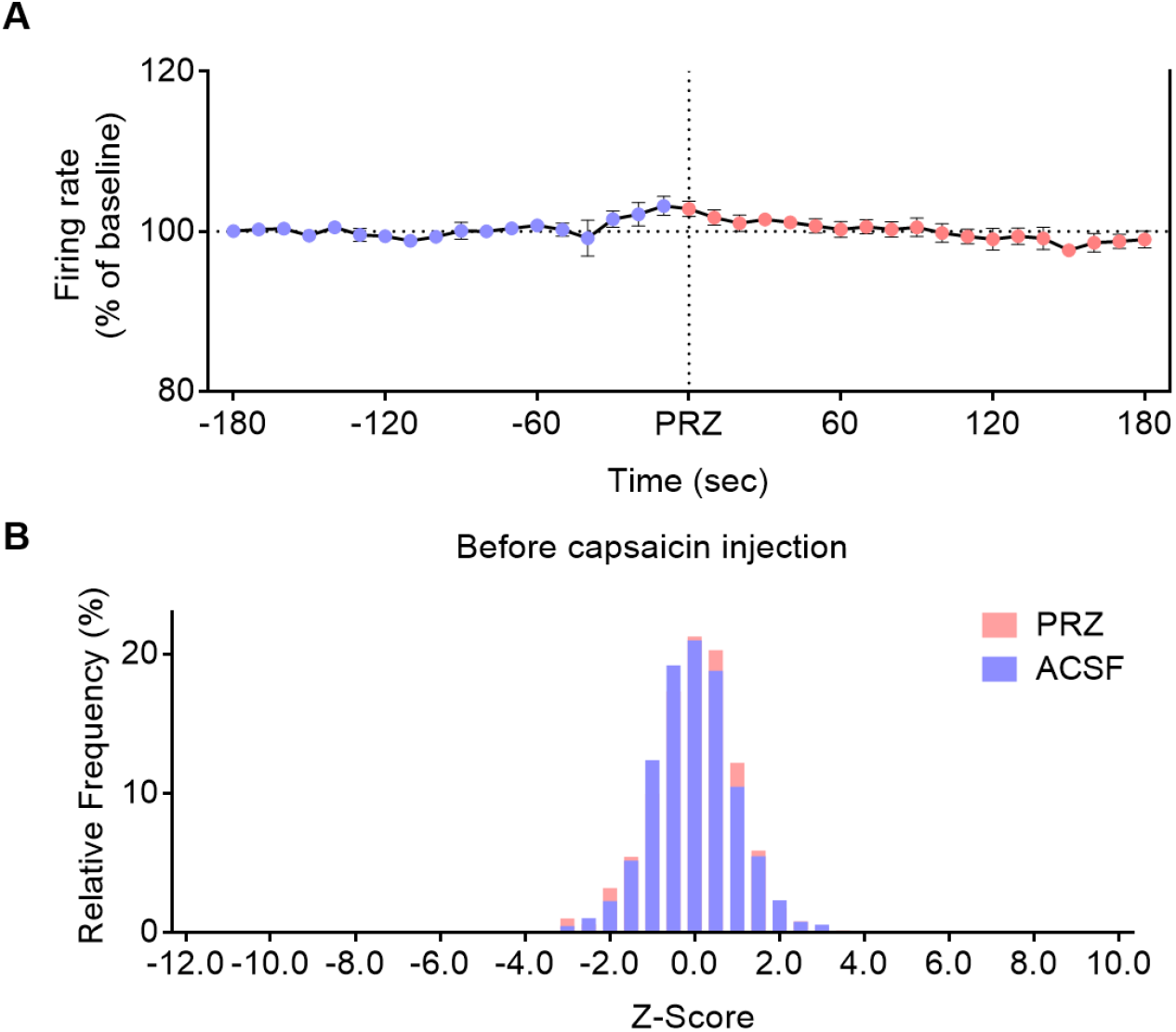
No change in PC firing rate through prazosin application. A, Plot showing PC spontaneous firing rate before (blue dots) and after prazosin application (red dots) (n = 6 slices in 3 mice). B, Bar graph showing the Z-score of PC spontaneous firing rate before (blue) and after prazosin application (red) in vivo.

**Fig. S5.**
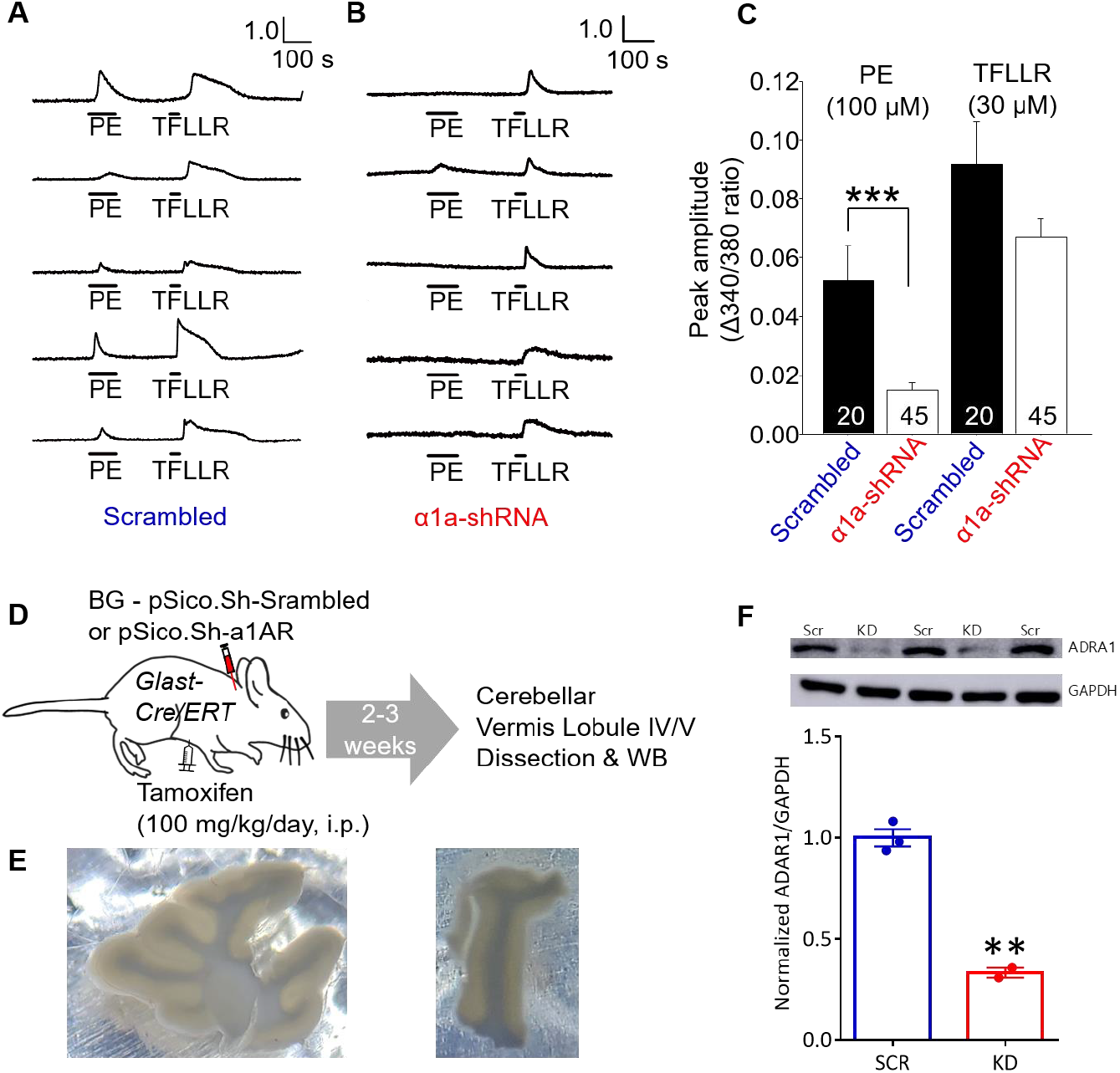
Validation of BG-specific genetic manipulation. (A-C) α1-AR KD development. A, Representative calcium traces of glial cells of Scrambled group by PE and TFLLR application. B, Representative calcium traces of glial cells of α1a-shRNA group by PE and TFLLR application. C, Summarizing bar graph showing calcium transient amplitude of glial cells (n = 20 glial cells in Scrambled group; n = 45 glial cells in α1a-shRNA group; PE 100 μM, ** p = 0.0376; TFLLR 30 μM, N.S; Two-way ANOVA). D, A diagram for experimental design. BG-specific α1 -AR KD virus was delivered and genetic knockdown was validated by western blot analysis. Cerebellar lobule 4/5 where the virus was delivered were selectively dissected. E, Example images of dissection of cerebellar lobule 4/5. F, Representative images (top) and its summarizing bar graph showing significant decrease of ADRA1 expression in KD compared to scrambled group. (Difference between groups = 0.67 ± 0.06; ** p = 0.0015; Unpaired t test).

**Fig. S6.**
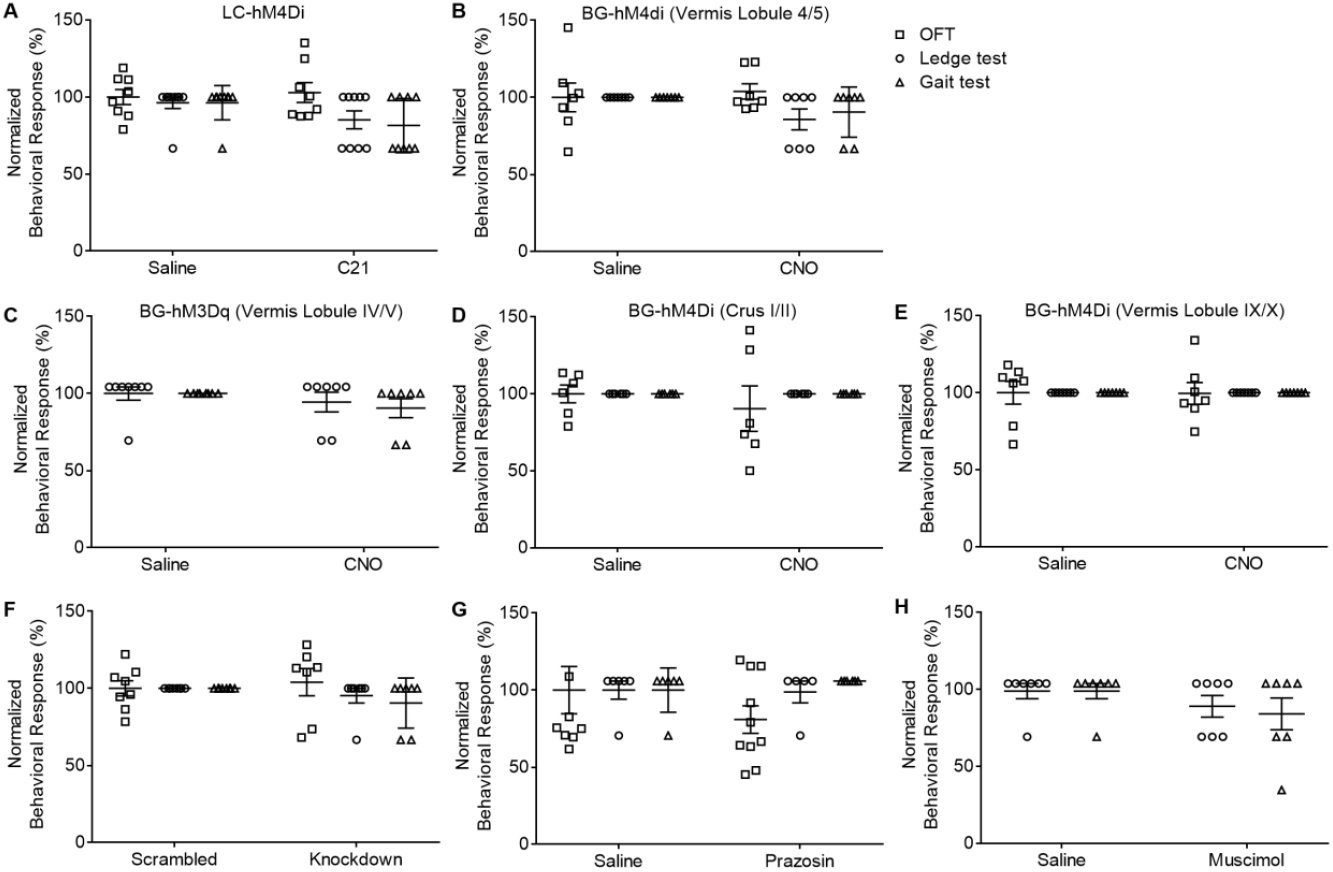
Locomotion was compatible among experimental groups. There was no significance in all experiments, and each group was as follows (Two-way ANOVA). A, C21 local application of LC hM4Di group (n = 9 mice saline or C21). B, CNO i.p. injection of BG hM4Di in cerebellar vermis lobule 4/5 group (n = 7 mice in saline or CNO group). C, CNO i.p. injection of BG hM3Dq in cerebellar vermis lobule 4/5 group (n = 8 mice in saline group; n = 7mice in CNO group). D, CNO i.p. injection of BG hM4Di in cerebellar Crus 1/2 group (n = 6 mice in saline or CNO group). E, CNO i.p. injection of BG hM4Di in cerebellar vermis lobule 9/10 group (n = 7 mice in saline or CNO group). F, BG specific α1-AR knockdown group (n = 8 mice in scrambled group; n = 7 mice in Knockdown group). G, Prazosin application in cerebellar vermis lobule 4/5 (n = 6 mice in saline or Prazosin group). H, Musimol application in DCN (n = 7 mice in saline or Muscimol group).

**Fig. S7.**
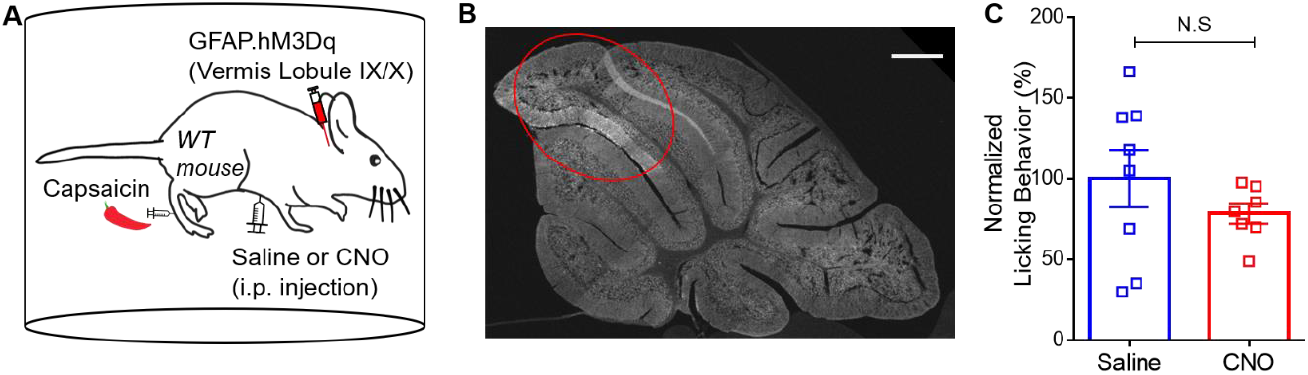
Capsaicin-induced nocifensive behavior during BG hM3Dq activation. **A,** A diagram for experimental design. Normalized percentage of capsaicin-induced licking behavior of saline and BG hM3Dq groups (Virus expression in cerebellar vermis lobule 4/5). **B,** Virus expression in cerebellar vermis lobule 4/5. **C,** Capsaicin-induced licking behavior in the presence of saline or CNO (n=9 mice for saline group; n=10 mice for CNO group; N.S., p=0.2985; Unpaired t-test). Scale bar, 500 μm.

**Fig. S8.**
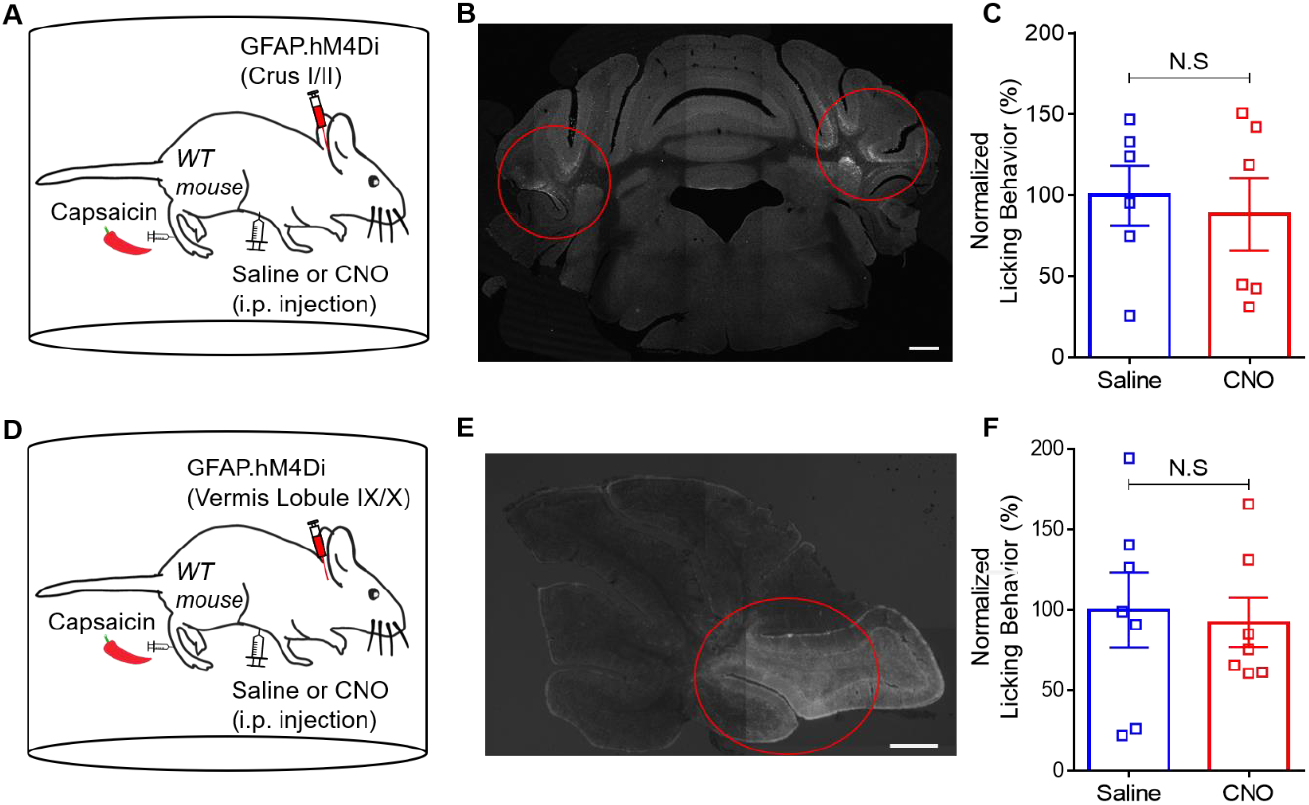
Capsaicin-induced nocifensive behavior during BG hM4Di activation in other cerebellar regions. A, A diagram for experimental design. Normalized percentage of capsaicin-induced licking behavior of saline and BG hM4Di groups (Virus expression in cerebellar crus 1/2). B, Virus expression in cerebellar crus 1/2. C, Capsaicin-induced licking behavior in the presence of saline or CNO (n = 6 mice for each group; N.S., p = 0.6985; Unpaired t-test). Scale bar, 500 μm. D, A diagram for experimental design. Normalized percentage of capsaicin-induced licking behavior of saline and BG hM4Di groups (Virus expression in cerebellar vermis lobule 9/10). E, Virus expression in cerebellar vermis lobule 9/10. F, Capsaicin-induced licking behavior in the presence of saline or CNO (n = 7 mice for each group; N.S., p = 0.7851; Unpaired t-test). Scale bar, 500 μm.

**Fig. S9.**
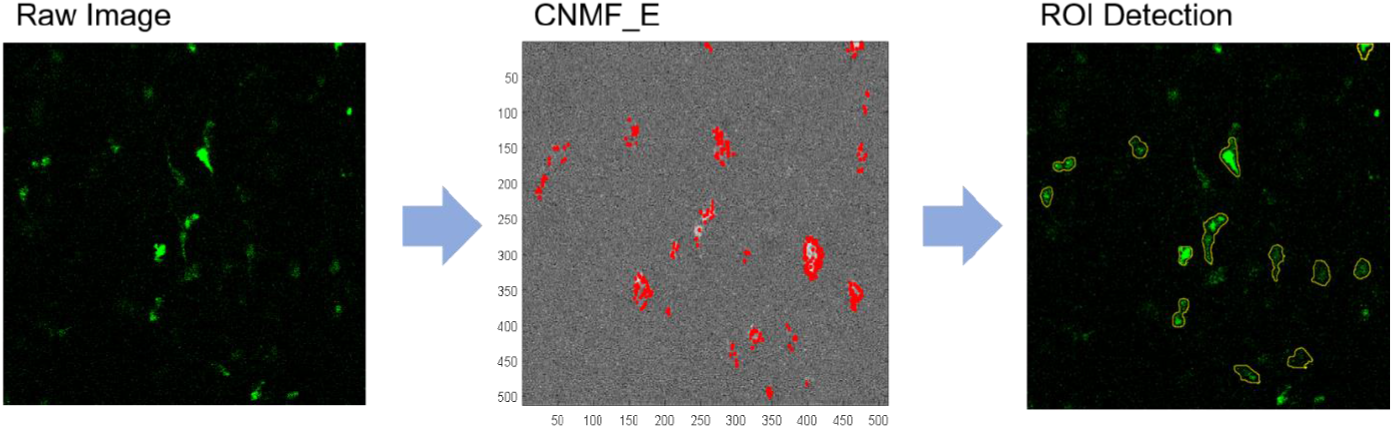
ROI detection of LC terminal using the CNMF_E.

